# LRP2 is a potential molecular target for pathological myopia

**DOI:** 10.1101/2025.03.04.641381

**Authors:** Kimberley Delaunay, Emilie Picard, Patricia Lassiaz, Laurent Jonet, Vidjea Cannaya, José Maria Ruiz Moreno, Kentaro Kojima, Henrik Vorum, Bent Honoré, Jorge R Medrano, Lasse Jørgensen Cehofski, Eric Pussard, Renata Kozyraki, Alicia Torriglia, Olivier Cases, Francine Behar-Cohen

## Abstract

High myopia (HM) and posterior staphyloma (PS) are major causes of vision loss worldwide. Genetic and environmental factors, especially light exposure, contribute to myopia. Mutations in low-density lipoprotein-related receptor type 2, LRP2 cause syndromic myopia, and the Foxg1-Cre-Lrp2lox/lox mouse is a model for myopia and PS but the involvement of LRP2 in non-syndromic HM (NSHM) was unknown. We showed that LRP2 levels were decreased in the vitreous of 25 patients with NSHM and PS, and that in human donor eyes affected by PS, LRP2 expression was reduced in the neural retina and retinal pigment epithelium (RPE). The morphologic changes of these RPE were similar to those observed in the RPE of the Foxg1- Cre-Lrp2lox/lox mouse. In iPSc-derived human RPE cells (iRPE), LRP2 was expressed at a functional location, and LRP2 silencing by a specific siRNA regulated genes belonging to pathways involved in eye and neuronal development, visual perception, tissue remodeling, hormone metabolism and RPE structure, demonstrating that LRP2 orchestrates in RPE, functions that are essential for eye growth. Exposure of iRPE to light with LEDs of different wavelengths upregulated LRP2 expression, with higher efficacy for red light. Conversely, LRP2 expression was downregulated after cortisol exposure. Our findings link LRP2 to myopization and environmental factors and highlight its role in NSHM and PS in humans. LRP2 appears to be a viable target for interventional strategies in the treatment of NSHM.

**One Sentence Summary:** LRP2, pivotal in the regulation of eye growth, exhibits a decrease in high myopic eyes. Its expression is augmented by light exposure in retinal pigment epithelial cells.

## INTRODUCTION

The prevalence of myopia, a refractive error commonly caused by abnormal elongation of the eyeball, is increasing rapidly worldwide, with the global prevalence expected to exceed 50% by 2050^1^. The refractive error can be corrected by optical or surgical means, but high myopia (HM), greater than 6 diopters, is often associated with sight-threatening retinal complications, including retinal detachment and maculopathy secondary to posterior staphyloma (PS), a focal protrusion of the ocular globe with an extreme thinning of the retinal pigment epithelium (RPE), the choroid, the neural retina, and the sclera^2,3^. Intensive research has led to the development of intervention strategies to reduce myopia progression, including atropine drops, defocus lenses or light therapy, which have significant but moderate effects^4^. To date, there is no method to prevent HM and its blinding complications, which is a major public health problem^5^.

Myopia results from a complex interplay between genetic and environmental factors^6,7^, the most widely accepted of which is exposure to daylight, to the point where light therapy has been proposed as a preventive strategy. However, the respective effects of different light components (such as luminance, spectral composition, temporal modulation) and of circadian regulations^8–10^ remain incompletely understood. Several hundreds of genes most of which are involved in TGF-beta signaling, collagen synthesis and retinal signal transduction have been associated with myopia^11^ and the study of monogenic syndromic myopia can help understanding of the mechanisms involved in multifactorial myopia. Variants in *LRP2 (*low-density lipoprotein receptor-related protein 2), which encodes megalin/LRP2 causes the rare Donnai-Barrow syndrome, which combines syndromic myopia and PS, hypertelorism, sensory neuronal loss, partial agenesis of the corpus callosum and proteinuria^12–14^. The mouse model conditionally knockout for *Lrp2* in ocular tissues (Foxg1-Cre-Lrp2lox/lox) developed postnatal myopia and a posterior protrusion of the globe very similar to myopic staphyloma^15^.

Furthermore, conditional mutants in which *Lrp2* has been knocked out only in the retinal pigment epithelium (RPE) also developed myopia and staphyloma^16^. In addition, the transcriptional downregulation of *Lrp2* in the RPE during the first three postnatal weeks promoted rapid eye growth and, high *Lrp2* expression in RPE of adult mice ensured that eye growth stopped at the correct size^17^. Taken together, these previous findings highlight the key role played by LRP2 in RPE-driven regulation of eye growth and in staphyloma formation. However, the mechanisms involved are still unclear.

LRP2 is a large transmembrane glycoprotein detected at the apical membrane of various epithelial cells where it participates in clathrin-mediated endocytosis, capturing and releasing ligands that either undergo degradation in lysosomes or are transported via a transcytosis route and secreted at the basal pole of the cell^18^. It regulates the extracellular concentrations of molecules, including vitamins and hormones ^19,20^ and modulates the activity of morphogens such as sonic hedgehog (SHH) or bone morphogenetic protein 4 (BMP4) during development^21,22^. Via endosomes recycling, LRP2 also controls cell shape and planar cell polarity ^23,24^. The availability of LRP2 at the cell surface is subjected to regulatory mechanisms, including the phosphorylation of its cytoplasmic domain, a PKC-regulated metallo-protease ectodomain shedding followed by the processing of the membrane bound C-terminal by gamma-secretase that produces a soluble intracellular domain and, exosome secretion^25^. In murine RPE, LRP2 is predominantly located at the apical plasma membrane, where it facilitates the uptake of albumin or transferrin^19^. But the mechanisms by which LRP2 expressed in the RPE controls eye growth are poorly understood. Whether LRP2 is also involved in NSHM is unknown.

We set out to investigate the involvement of LRP2 in NSHM in humans. We used human RPE derived from iPSc to study the transcriptional consequences of LRP2 silencing and the effect of environmental factors, including light and cortisol exposure, on LRP2 expression.

## Results

### Levels of LRP2 are decreased in the vitreous of NSHM eyes with PS

LRP2 levels was evaluated in vitreous samples of high myopic patients with PS who underwent pars plana vitrectomy for myopic tractional maculopathy, macular hole or epiretinal membrane and compared to vitreous from emmetropic patients operated for epiretinal membrane or for intraocular lens luxation. These measurements were carried out on a Japanese population and on a European Caucasian population by two independent research groups and two different analysis methods.

A proteomic analysis was performed on the vitreous from 15 NSHM Japanese patients compared to 10 controls. **Table 1** recapitulates the demographic characteristics and surgical indications. Axial length in the HM group was 29.33±1.94mm and all patients had posterior staphyloma. Proteomic analysis of the vitreous performed by label-free quantitative liquid chromatography-tandem mass spectrometry identified 332 proteins (**Data File S1**) of which were 73 differentially expressed proteins (DEPs), including LRP2 (fold change = 0.5; *p* = 0.016) (**Fig. 1A**; **Table 2**). The top up-regulated DEPs included apolipoprotein D, transforming growth factor-beta-induced ig-H3 (TGFBI), complement factor D and thrombospondin-4. On the other hand, versican core protein, S-arrestin, collagen IX alpha-2 chain, LRP2, and retinol-binding protein 3 (IRBP) were reduced. The most significant biological processes associated with downregulated proteins were linked to nervous system development and negative regulation of proteolysis **(fig. S1)**. Molecular functions related to the significantly regulated proteins included reduced low-density lipoprotein binding, calcium ion binding, and chaperone binding, whereas serine-type endopeptidase inhibitor activity and integrin binding were elevated **(fig. S1).** The down-regulated proteins were predominantly located in the extracellular region and extracellular space, while up-regulated proteins were mostly linked to extracellular matrix (**fig. S1**). Abnormal retinal morphology, myopia, and retinal detachment were the most significant phenotypes associated with the differentially expressed proteins (**Fig. 1B**). The most significant connected protein cluster contained 14 proteins, including LRP2 and its ligands clusterin (CLU) and APOE (**Fig. 1C** and **fig. S2A**), which were enriched in neurodegenerative disorders (Gene Ontology Disease, **fig. S2B**).

**Fig. 1.**
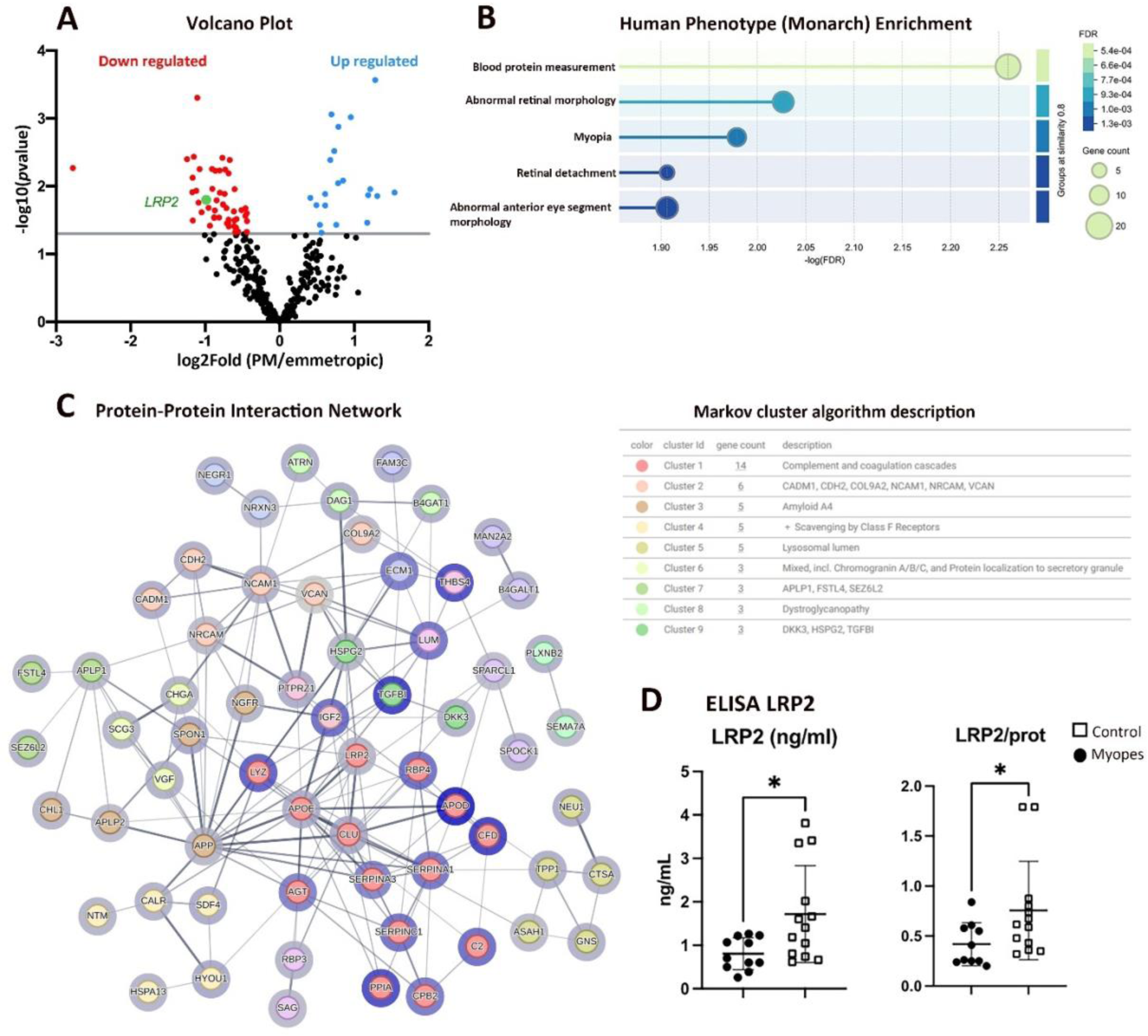
Analysis of proteins in the vitreous of myopic and emmetropic patients. (**A**) Volcano plot of differentially expressed proteins. The rightmost part of the plot (blue circles) showed the 20 up-regulated proteins and the leftmost (red circles) the 53 down regulated proteins. (**B**) Monarch analysis enrichment of down regulated proteins, only the top 5 terms were displayed. Gene count associated with a particular Monarch term is indicated. (**C**) Protein- Protein Interactions (PPI) using STRING analysis displayed the signaling network between the differentially expressed proteins. Nodes in blue circles refer to upregulated proteins, nodes in grey circles refer to down regulated proteins. Markov cluster algorithm analysis indicated the main interactomes. The nine top interactomes are indicated at the right. Protein names are provided in Table 1. (**D**) Concentrations of LRP2 protein in ng/ml in undiluted vitreous from control (n = 13) and NSHM myopic (n = 10) eyes quantified with ELISA. The graph on the right showed the ration of LRP2 compared to the total protein content. Statistical analysis was performed using the Mann-Whitney test, \**p*<0.05.

**Table 1.** Summary of human vitreous samples.

| Characteristics | Mass spectrometry analysis |  | ELISA |  |
| --- | --- | --- | --- | --- |
|  | myopia (n=15) | Controls (n=10) | myopia(n=10) | Controls (n=13) |
| Gender (male/female) | 3/12 | 4/6 | 5/5 | 8/5 |
| Age (year) | 71±8.6 | 71.6±8.5 | 63±4 | 67.3±18 |
| Axial length (mm) | 29.33±1.94 | 24.11±1.57 | 29.67±2 | 22.6±0.7 |
| Posterior Staphyloma | 15 | 0 | 10 | 0 |
| **<u>Indications for surgery</u>** |  |  |  |  |
| Epiretinal membrane | 4 | 10 | 2 | 10 |
| Myopic Tractional maculopathy | 11 |  | 8 |  |
| Macular hole | 1 |  |  | 1 |
| Intraocular lens luxation |  |  |  | 2 |

**Table 2.**
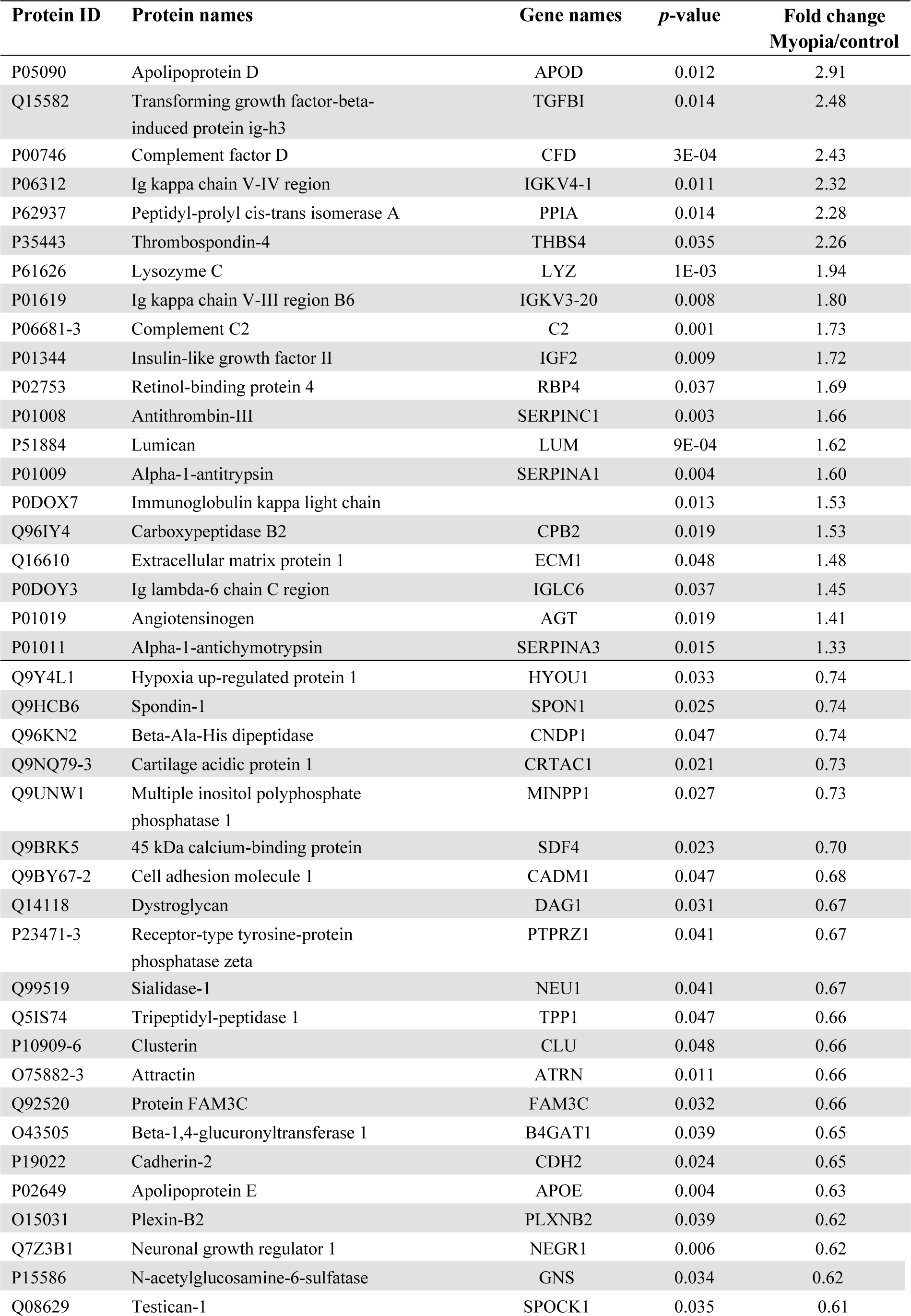
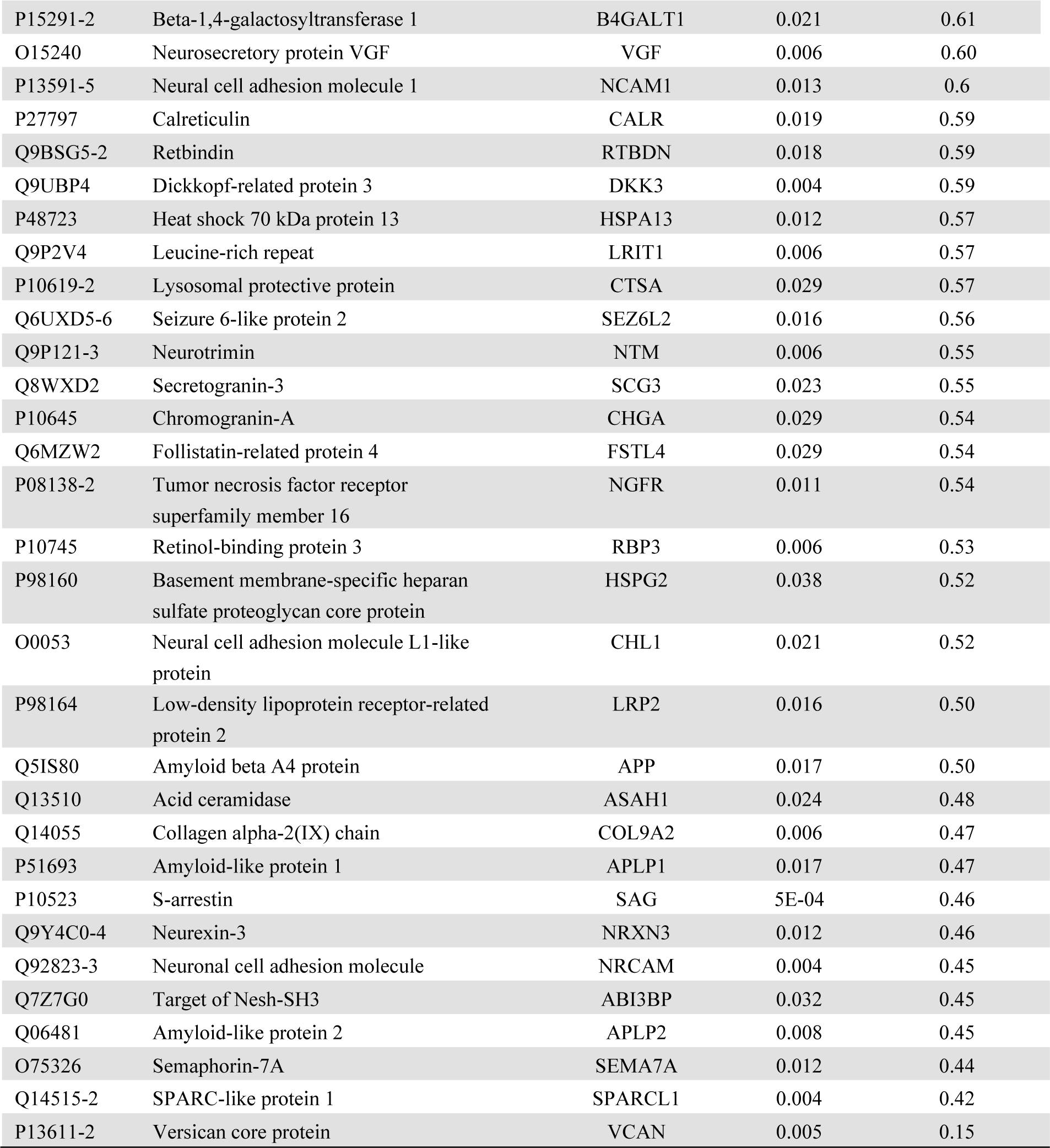
Significantly Differentially Expressed Proteins in High Myopia compared to Controls.

In the European Caucasian group of patients, 10 HM patients and 13 emmetropic patients were included. Table 1 recapitulates the demographic characteristics and surgical indication. In the HM group, axial length was 29.67±2 mm and all eyes had posterior staphyloma. LRP2 was measured by enzyme-linked immunosorbent assay (ELISA) analysis on vitreous samples. LRP2 levels and LRP2/protein were significantly lower in HM eyes (0.8 ± 0.36 ng/ml and 0.41±0.21) than in emmetropic eyes (1.7 ± 1.1 ng/ml and 0.75±0.49) (*p*=0.01 and 0.02 respectively) (**ig. 1D**).

### Pathological features of high myopic human eyes with posterior staphyloma

Two highly myopic post-mortem eyes, with axial lengths of 32 mm and 31 mm for the right and left eye, and a posterior staphyloma, were studied. The pre-mortem spectral domain optical coherence tomography (SD-OCT) images of the right pseudophakic HM eye revealed posterior incurvation of the PS with thinning of the retina and the choroid (**Fig. 2, A andB**, black arrows), complicated by myopic tractional maculopathy (MTM) with foveoschisis (**Fig. 2, A and B**; stars and white stars). The SD-OCT images of the left eye were of low quality due to corneal opacification. On the post-mortem enucleated donor eyeball, the PS with scleral thinning was clearly visible (**Fig. 2C**). Peripapillary atrophy was visible on macroscopic pictures of the open eyeball (**Fig. 2, D and E**), and the PS was located temporal to the fovea (**Fig. 2, D and E;** black arrow). A cross-section at the level of the PS (**Fig. 2G**) showed disorganization of the retinal layers and cysts (arrows) and extreme retinal thinning at the bottom of the PS (**Fig. 2G**; double arrow) compared to emmetropic retina (**Fig. 2F**). RPE cells outside the PS showed clear disorganization (**Fig. 2, H to J**), with mislocalization of melanosomes and aggregations of pigments. More precise cellular disorders were characterized using markers of different cell types on immunohistochemistry of emmetropic and HM human retina cryosections (**Fig. 3**). In the control retina, glial Müller cells (RMG) identified by glutamine synthetase expression (GS) extended their processes from the inner limiting membrane (ILM) to the outer limiting membrane (OLM) (**Fig. 3, A and B**). In the HM retina, RMG processes were disorganized and lost their alignments within the thinned layers (**Fig. 3C**). The RMG processes organized around cysts (**Fig. 3D**) and invaded the bottom of the PS that was covered by RMG cells (**Fig. 3E**). The neuronal-specific marker, tubulin beta 3, labeled the nerve fiber layer, the ganglion cells (RGC) and their extensions, and amacrine cells (**Fig. 3, F and G**). In the HM retina, a severe thinning of the nerve fiber layer (**Fig. 3H**) was observed together with a reduction of RGC (**Fig. 3, H and I**). Blue and green cones (**Fig. 3, J and K**), labeled by opsins, showed significant disorganization in HM retina with greater loss of blue cones (**Fig. 3, L and M**). Finally, the astrocytic-specific marker GFAP, which stains astrocytes and the end feet of RMG cells in the emmetropic retina (**Fig. 3, N and O**), showed a dense layer at the surface of HM retina, which could correspond to the vitreoretinal tractional membrane seen on SD-OCT (**Fig. 2, A and B**) but the RMG invading the posterior staphyloma did not express GFAP. The retina of the HM eye exhibited loss of ganglion cells and cones, as well as thinning of all retinal layers, and cystic degeneration accompanied by glial Müller cells disorganization. Extreme degeneration was present in the PS area which was filled by glial Müller cells.

**Fig. 2.**
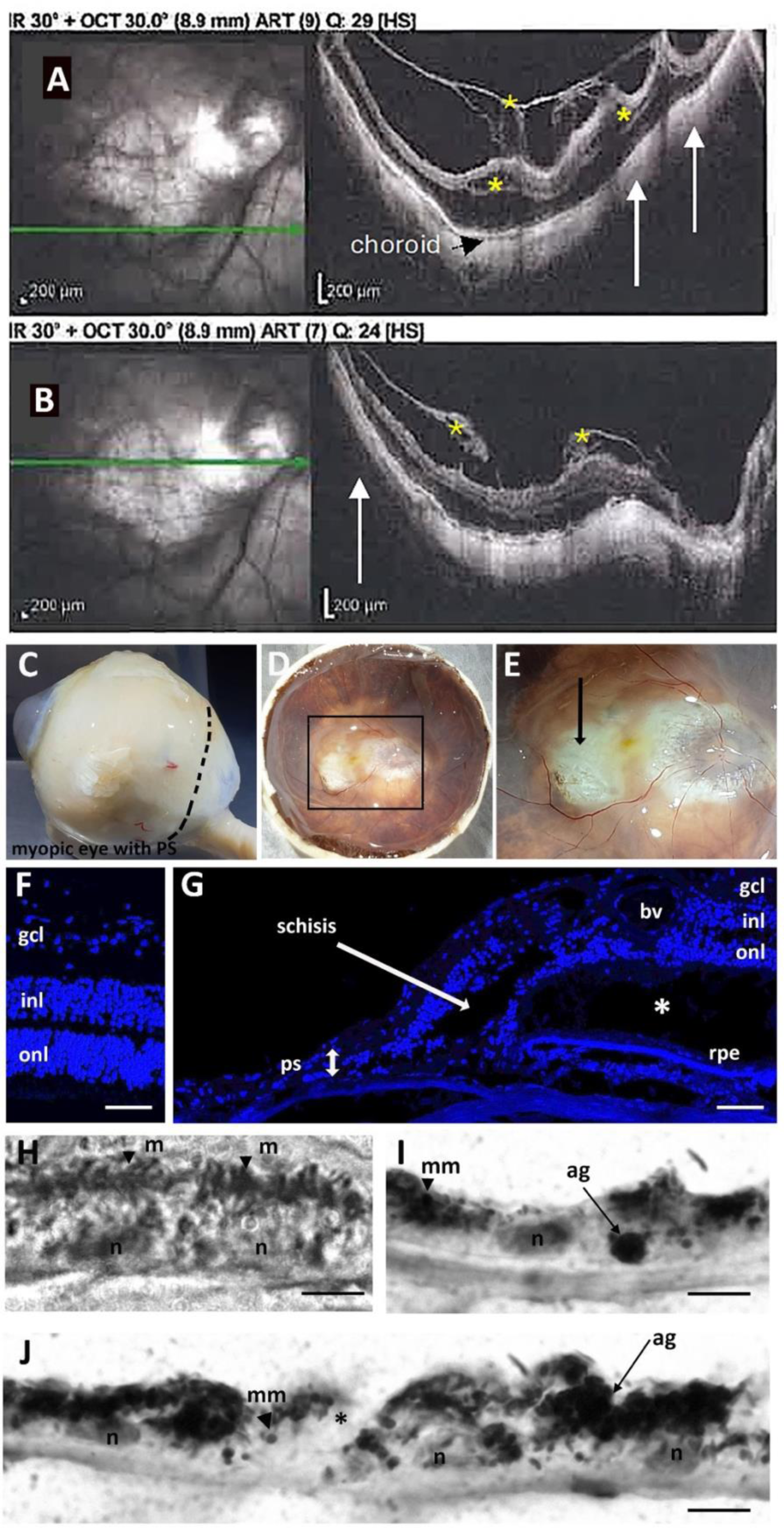
Ophthalmologic imaging and histological analysis of the right eye of a HM donor. **(A-B)** Pre mortem SD-OCT scans of the right eye with inferior cross-section (**A**) and macular cross-section (**B**), showing the PS (white arrows) and tractional maculopathy (yellow stars). The myopic eye displayed an atrophy of the macula delineated by small arrows and the posterior incurvation indicated the uncomplicated PS. **(C)** Gross morphology of a the high myopic right enucleated post mortem eye with a posterior staphyloma (PS). (**D**) Macroscopic image of the posterior chamber after dissection showing the posterior staphyloma (ps). (**E**) Insert box of (**D**) focusing on fovea and ps (black arrow). **(F**) DAPI-stained section of a normal human retina near the macula. (**G**) DAPI-stained section of the high myopic retina with a PS (double headed arrow) shows a severe reduction of thickness of all retinal layers, adjacent to the PS. Schisis (white arrow) is observed between the inner nuclear layer (inl) and the outer nuclear layer (onl). A cyst is observed between the RPE and the outer segments layer (asterisk). bv: blood vessel. (**H**) RPE cells in human retina form a single layer of pigmented cells as observed on light transmission microscopy. Melanosomes (m) have an elongated shape and nuclei (n) are spaced at regular intervals. (**I**, **J**) In the HM donor eye, RPE cells are still forming a single layer. RPE cells are abnormally large, their apical domain is reduced, and melanosomes aggregate in macromelanosomes (mm) or macrostructures (ag). Scale bars: F,G = 120 μm; H = 75 μm; H-J = 5 μm.

**Fig 3.**
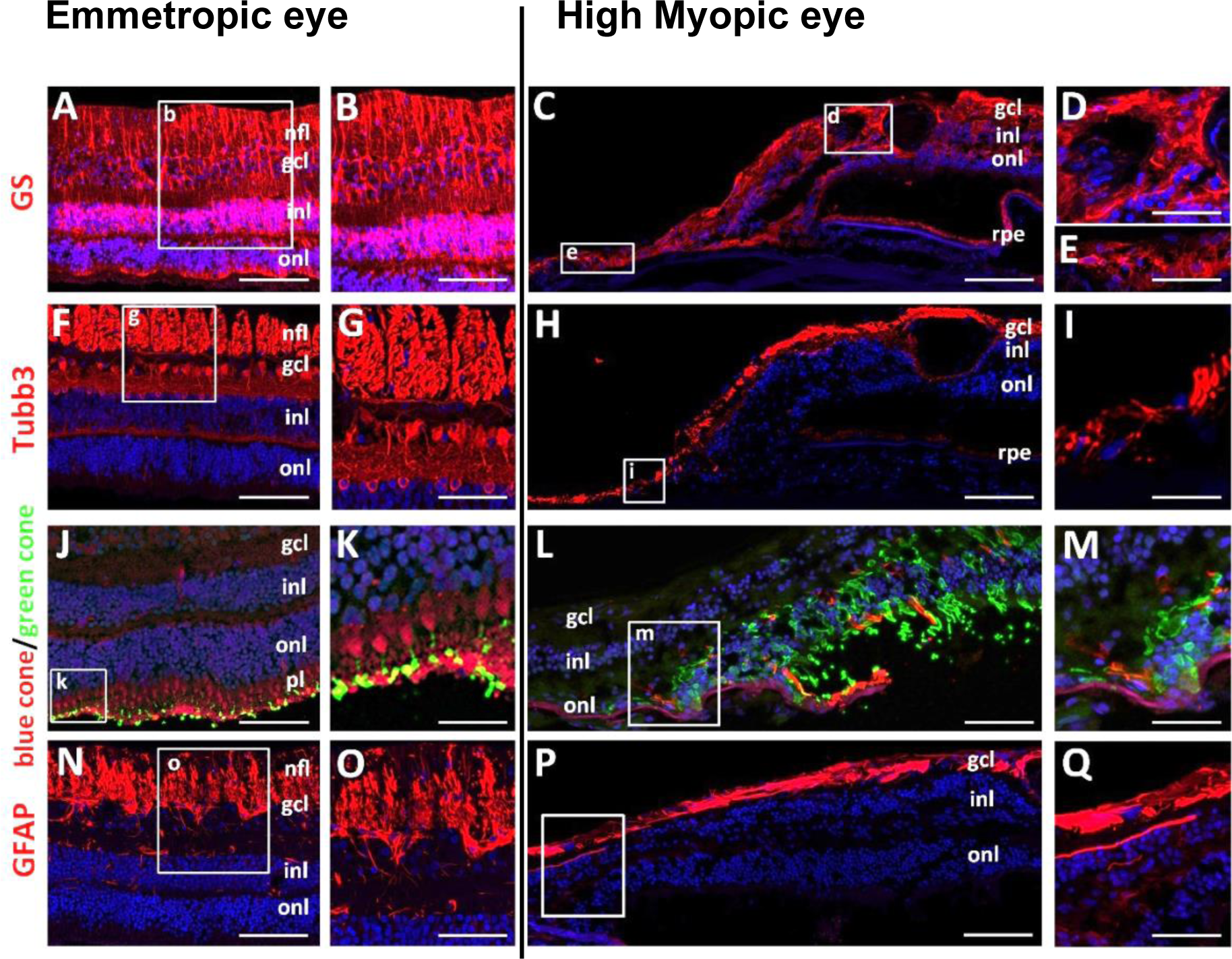
Immunolabelling of retinal cells in the emmetropic and HM retina. (**A, B**) Glutamine synthetase (GS) labels Müller glial cells in the emmetropic retina, extending radially from the nerve fiber layer (nfl) to the onl (**C, D**) In HM retina, GS-positive cells extend in all directions and thicken at the retinal surface. At higher magnification, GS-positive cells surround a cyst in the ganglion cell layer (gcl) (**C**) and fill the bottom of the staphyloma where almost no retina remains **(E)**. (**F, G**) Tubulin beta-3 (Tubb3) is a neuronal marker of retinal ganglion (RGC) and amacrine cells in the emmetropic retina. (**H**) In HM retina, the number of Tubb3-amacrine cells and RGCs is reduced. Higher magnification shows Tubb3-positive axons browsing at the surface of the staphyloma. (**J, K**) Blue (red staining) and green (green staining) opsin markers reveal the distribution of blue and green cones in the emmetropic retina. (**L**, **M**) In HM retina, the number of blue cones is reduced (**L**) and high magnification shows the accumulation of green opsin in abnormally shaped outer segments. (**N, O**) Glial fibrillary acidic protein (GFAP) labels astrocytes and glial Müller cells endfeet in the emmetropic retina. (**P**, **Q**) In the HM retina, the GFAP-positive endfeet are reduced. (**Q**) At higher magnification, a GFAP- positive membrane lays on the retinal surface but no positive cells are observed in the retina . Scale bars: A,F,J,N = 200 μm; B,G,O = 80 μm; K = 20 μm; C,H,L,P = 250 μm; D,E,I,M,Q = 60 μm.

### Decreased LRP2 in human HM with PS retina is associated with severe RPE disorganization similar to Foxg1-Cre-Lrp2lox/lox mouse RPE

In emmetropic human neural retina, LRP2 was expressed in RMG cells, co-localized with GS (**Fig. 4, A and B**). In the HM neural retina, significant disorganization of the retinal layers was evident at the margin of the staphyloma, where glial Müller cells spanning through layers and the subretinal space expressed low levels of LRP2 (**Fig. 4, C and D**). In the RPE cells, LRP2 expression was decreased in the HM compared to the emmetropic eye, as seen in both transversal sections from left eye (**Fig.e 4E**) and flat-mounts of the right eye (**Fig. 4, F to H**). In emmetropic RPE, LRP2 was located at the apical membrane and in vesicles (**Fig. 4E**; arrowhead), but also showed diffuse labeling underneath the Bruch membrane (BM), in the most inner part of the choroid where it stained the pillars of the choriocapillaris (**Fig. 4E**). In RPE flat-mounts of the emmetropic control eye, LRP2 was localized in punctiform sub- membrane clusters at the apical and lateral borders of the RPE cells (**Fig. 4, F and G**), at the cell membrane (**Fig. 4G**), and in clathrin-positive vesicles (**Fig. 4H**). On transversal sections of HM RPE cells (**Fig. 4I**), LRP2 was faintly detected (**Fig. 4, J to L**), at the cell membrane (**Fig. 4K**) and in very few clathrin-positive vesicles (**Fig. 4L**) with staining of LRP2 aggregated at the apical side (**Fig. 4I**; arrowhead) without any signal underneath BM (**Fig. 4I**). Along with LRP2 reduction, a parallel reduction in clathrin-positive vesicles (**Fig. 4L**) was observed. In HM eyes with PS, LRP2 was thus decreased not only in the vitreous but also in retina and RPE. RPE morphology was characterized by phalloidin staining that labels actin cytoskeleton. As opposed to the regular arrangement of RPE cells in emmetropic human eye (**Fig. 5A**), in HM eye, RPE displayed irregular organization, cell size dispersity (**Fig. 5B**) and doubling or breaks of the cell junctions (**Fig. 5, C and D**). At the bottom of the staphyloma, the extreme thinning and eye incurvation did not allow for RPE flat mounting.

**Fig. 4.**
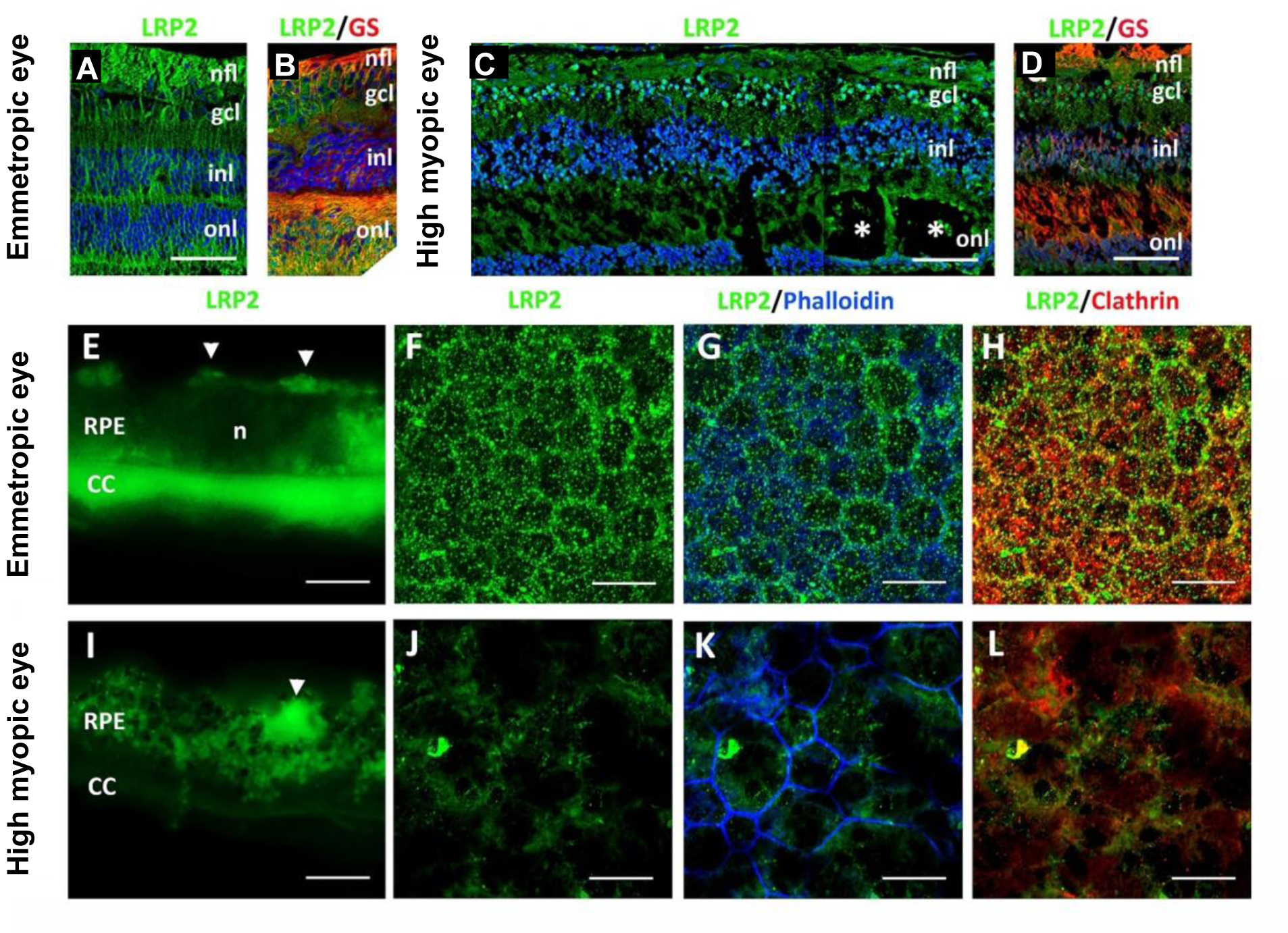
LRP2 immunolabelling in the emmetropic and HM retina. (**A,B**) In the emmetropic neural retina, LRP2 is expressed around RGCs and along GS positive glial Müller cells. (**C, D**) In the HM neural retina, LRP2 expression is greatly reduced in GS- positive cells, that surround cysts (stars). (**E**) In transversal sections of RPE cells, LRP2 is located at the apical pole (white arrowheads), in intracellular vesicles and at the basal pole of the RPE, and in the pillars of the choriocapillaris. (F, G) In RPE flat mounted preparation from the left emmetropic eye, LRP2 is distributed in cytoplasmic vesicles (**F**) along apical and lateral membranes (**G**) and most LRP2-positive vesicles are also positive for clathrin. (**I**) In transversal section of the HM RPE, LRP2 expression is greatly diminished and absent in the choriocapillaris. (**J to L**) In RPE flat mounted from the HM eye, LRP2 distribution is sparse and diffuse, and does not colocalize with clathrin (**L**) that is also diminished. Scale bars: A-D 200 μm; G-I, K,L 50 μm.

**Fig 5.**
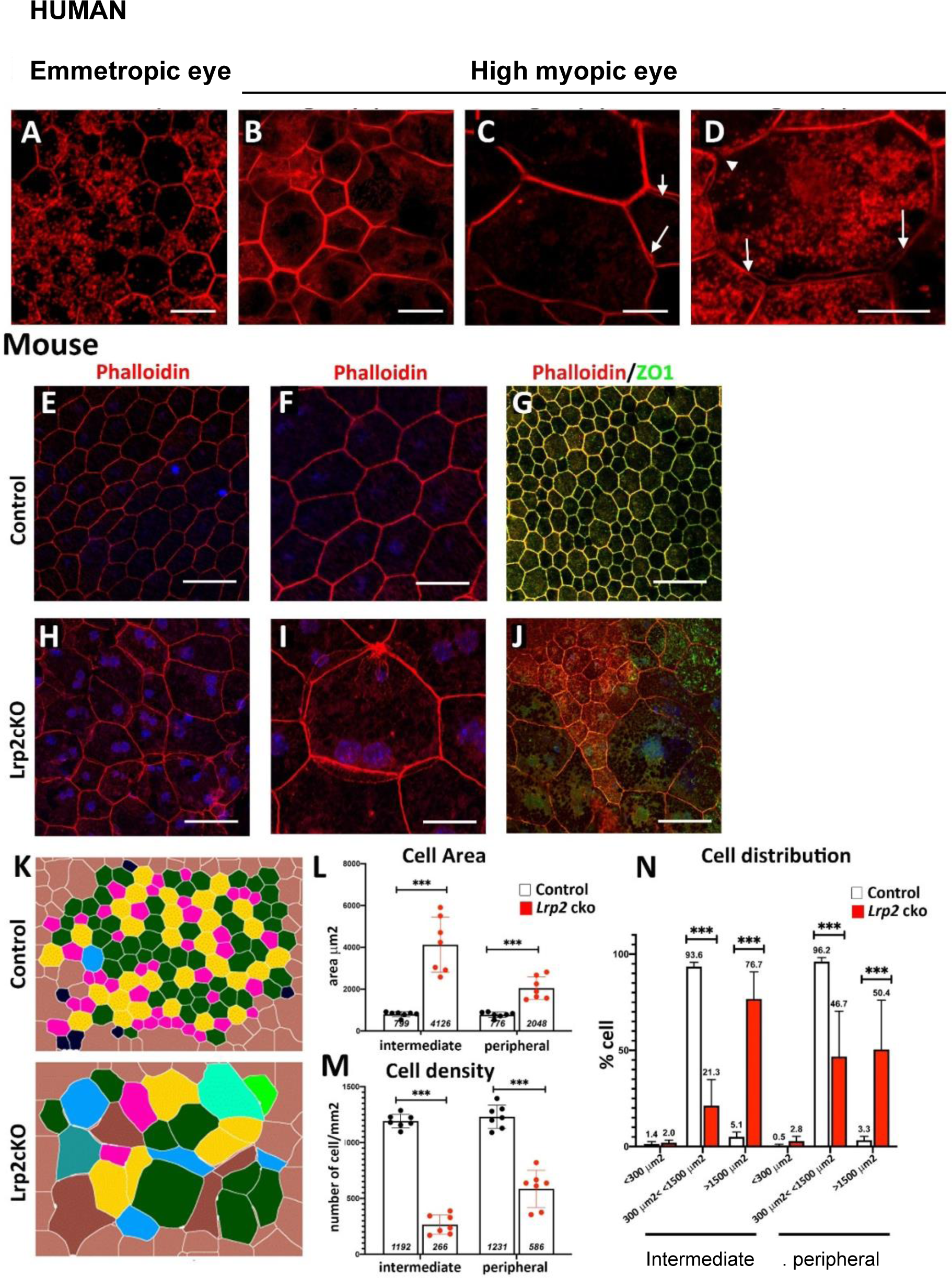
Organization of the RPE is altered in HM eye and in Lrp2cKO mouse**. (A)** Phalloidin staining reveals the geometric paving pattern of RPE in the emmetropic eye. (**B to D**) In HM eye, RPE geometric paving pattern is lost as most cells increase in size. Bicellular junctions display secondary actin arcs (**C**, arrows) and cell junctions are disorganized (**D** arrows). (**E, F**) Phalloidin staining on RPE flat-mount of WT control mouse shows the regular pattern of RPE cells. (**G**) Zonula Occludens 1 (ZO1) follows in the apical pole the distribution of phalloidin. (**H**, **I**) In Lrp2cKO RPE, the pseudo-geometric paving pattern is lost and is replaced by a tangle of cells whose surface has increased. (**J**) ZO1 is redistributed in the cytoplasm of abnormally shaped RPE cell. (**K**) Digital overlay reconstruction of control and, Lrp2cKO RPE indicates the increase size of RPE cells both at the periphery and at the intermediate level of the retina in Lrp2cKO. (**L**) Cell area in μm^2^, (**M**) cell density in number of cells/mm^2^, and (**N**) distribution of cells by sizes in the intermediate and peripheral retina of control compared to Lrp2cKO mice. Values represent the mean of cell size average of each sample (n=7) per genotype ± SEM. Mann-Whitney U test. *** *p* <0.001. Scale bars: A-D,E,H = 50 μm; F,I = 25 μm; G,J = 75 μm.

The RPE from Foxg1-Cre-Lrp2lox/lox mouse was studied to explore the impact of suppressed LRP2 expression on the RPE morphology. As opposed to control WT mice (**Fig. 5, E to F),** that showed the regular octagonal RPE cell surrounded by eight regular hexagonal cells, the transgenic mouse displayed irregularly arranged RPE with dramatic changes in cellular size and shape mostly posterior to the equator (**Fig. 5, H and I**). Membrane ZO-1 staining tight junction was lost and replaced by diffuse cytoplasmic staining (**Fig. 5J**) . At the periphery, areas with normal-sized RPE cells showed actin stress fibers (**Fig. 5J**). Automatic quantification (**Fig. 5K**) revealed that RPE cell size increased in the middle and peripheral retina (**Fig. 5, L and M**) and that cell density decreased mostly posterior to the equator (**Fig. 5N**).

The RPE from individuals with HM and PS, which exhibit a marked decrease in LRP2 expression, displayed morphological changes analogous to those observed in *Lrp2* invalidated mice.

### Transcriptional consequences of *LRP2* silencing in iRPE cells

To decipher the consequence of LRP2 downregulation in RPE cells and its potential link with myopia and PS, we used RNA interference to silence *LRP2* mRNA in iRPE cells (shLRP2 iRPE). We first evaluated the ability of differentiated iRPE to express LRP2 and other endocytosis markers. LRP2 was localized at the cell membrane from the apical to the basolateral side and in subapical vesicles (**Fig. 6A**), co-localizing at the apical pole with the receptor-mediated endocytic protein clathrin (**Fig. 6B**), partially with the early endocytic marker EEA1, (**Fig. 6C**) and with LAMP1, a membrane protein principally located in lysosomes (**Fig. 6D**). LRP2 being expressed in an appropriate spatial and functional location, iRPE cells were used to study the molecular consequences of LRP2 down-regulation.

**Fig. 6.**
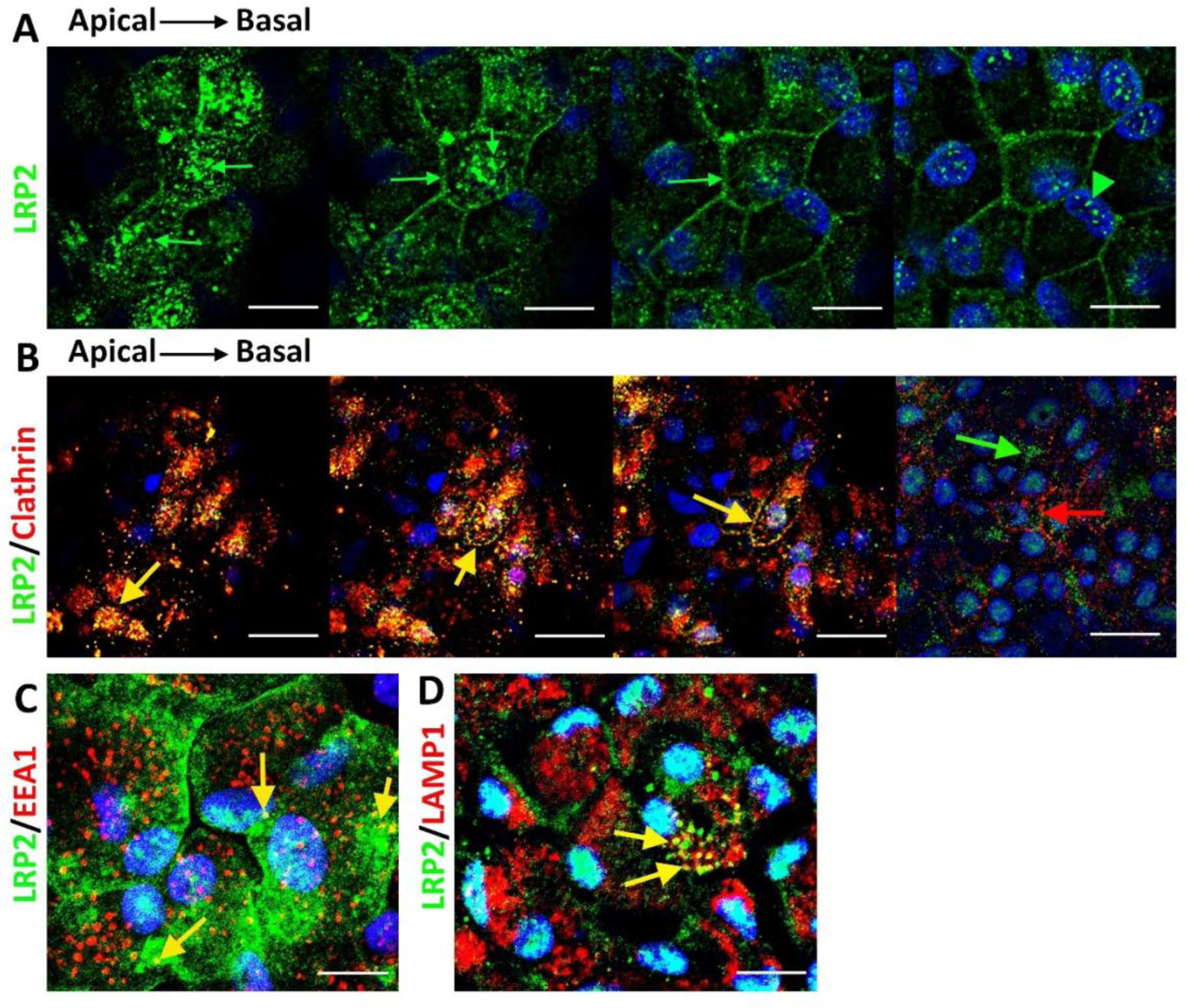
LRP2 localization in healthy human iRPE. (A) Consecutives confocal images from an apical to basal stack showing LRP2 accumulation in large vesicular structures in the apical pole (arrows), faintl localization at lateral membranes and in small cytoplasmic vesicles and in larger perinuclear vesicles (arrowheads). (B) Consecutives confocal images from an apical to basal stack show that LRP2 colocalized with the major protein of coated pits and apical endocytic vesicles, clathrin (yellow arrow). In the basal part of the cell, LRP2 (green arrow) did not colocalize with clathrin (red arrow). (C) Confocal image showing that LRP2 (green) partially colocalized with the endocytic marker, early endocytose antigen-1 (EEA1; red) (yellow arrow). (**D**) In the basal compartment, LRP2 (green) colocalize with LAMP1 (red) a specific lysosomal marker (yellow arrows). Scale bars: A,C,D = 10 μm; B = μm; C = 30 μm.

Bulk transcriptomes showed that shLRP2 specifically and efficiently reduced LRP2 mRNA expression by 40% after 48 hours, which was confirmed by quantitative PCR (**Fig. 7A**) and by western blot (**Fig. 7B**). Principal component analysis (PCA) showed the overall variation among samples, with a clear separation in the first three components between shLRP2 and scramble iRPE (**fig. S3**). We identified 118 differentially expressed transcripts between shLRP2 iRPE and scramble iRPE, including 75 down-regulated and 42 up-regulated transcripts, as shown in Volcano maps (**Fig. 7, C and D**; **Data File S2**). A list of differentially expressed genes (DEGs) along with their fold change and additional information about their roles in eye physiology, is provided in **Table 3** and **Table 4**. As expected, LRP2 was one of the most down- regulated genes by adjusted p-value. Along with LRP2, we identified several other RPE-specific apical membrane DEGs that were downregulated in shLRP2 iRPE, whilst upregulated DEGs were involved in the control of the circadian clock, cell cycle and in the regulation of TGF beta activity. The results of the KEGG enrichment analysis demonstrated that DEGs were mainly enriched in “retinol metabolism” (downregulation) and in “ECM receptor interaction” (upregulation) (**fig. S4A** and **Data File S3**). The GO-based enrichment analysis performed under the three categories: Biological Processes (BP), Cellular Components (CC), and Molecular Functions (MF) (**fig. S4, B to D** show the topmost enriched GO terms; **Table S3** has the complete list) identified the downregulation of genes related to molecular transmembrane transporter activity, fat soluble vitamin metabolic process (related to retinoic acid and vitamin D), ion homeostasis (**Fig. 7E**; **fig. S4, B and D**) and apical and basal plasma membranes (**fig. S4C** and **fig. S5A**). The upregulated DEGs were enriched in GO terms related to extracellular matrix and cytoskeleton arrangement (**fig. S4, C and D**). Using reactome pathways analysis (**Table S3**); the top downregulated proteins included the canonical retinoid cycle in vision and solute carrier-mediated transmembrane transport (**fig. S4E**), while the top upregulated DEGs included kinesins, collagen biosynthesis and modifying enzymes (**fig. S4C**). Human phenotype ontology (HPO) enrichment analysis (**Data File S3**) identified chorioretinal atrophy, RPE atrophy, and progressive night blindness (**fig. S4F**). When performing an enrichment analysis with LRP2 as a common factor (**Data File S4**), enriched GO terms were related to the apical plasma membrane, solute transmembrane transporters, fat soluble vitamin metabolic processes, sensory perception, and steroid metabolic processes (**Fig. 7F**). Upregulated enriched GO terms were related to the cytoskeleton and collagen-containing extracellular matrix (**Data File S4**). Reactome pathway analysis identified three downregulated pathways: visual phototransduction, metabolism of steroids, and sensory perception (**Fig. 7G**). **Table 5** recapitulates selected gene sets associated with LRP2 reduction in iRPE cells and their potential link with myopia and PS development.

**Fig. 7.**
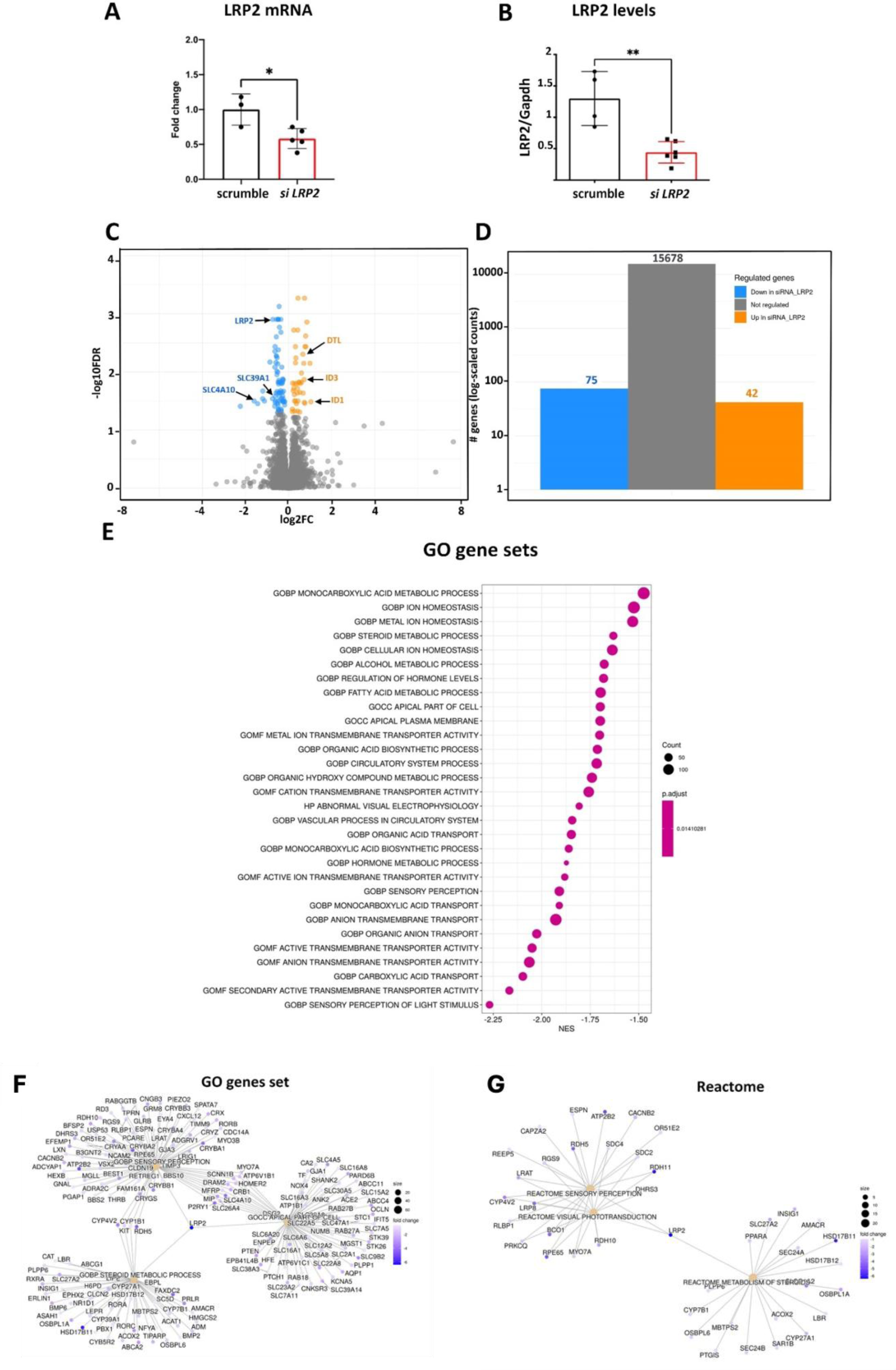
Differentially expressed genes (DEGs) between *shLRP2* and *scrambled* iRPE cells. (A) *Lrp2* mRNA quantification between *scrambled* (n = 4) and *shLRP2* (n = 4) iRPE (\**p*<0.05). (B) LRP2 levels as expressed by the ratio LRP2/GAPDH between *scrambled* (n = 4) and *shLRP2* (n = 4) iRPE (\*\**p*<0.01). (**C**) Volcano plot of DEGs between *shLRP2* and *scrambled* iRPE cells with –Log10 of the adjusted *p* value on the Y-axis and Log2 of the fold change expression on the X-axis. Some genes including LRP2 are indicated. (**D**) Graph bars show the number of downregulated (75), upregulated (42) and unchanged (15678) genes (**E**). The top 30 most enriched GO terms (selected based on the *p*-values) of regulated genes. Analysis was performed using Metascape.Category netplot gene enrichment analysis considering only LRP2 as a common factor with 3 GO terms, sensory perception, import across plasma membrane and steroid metabolic processes (**F**) and with 2 Reactome terms, sensory perception, visual transduction and metabolism (**G**). Fold change (color codes on each graph) for each gene of the selected GO term is indicated. Analysis was performed using Metascape. Absolute normalized enrichment allowed to identify terms that were upregulated or downregulated using all differentially expressed genes.

**Table 3:**
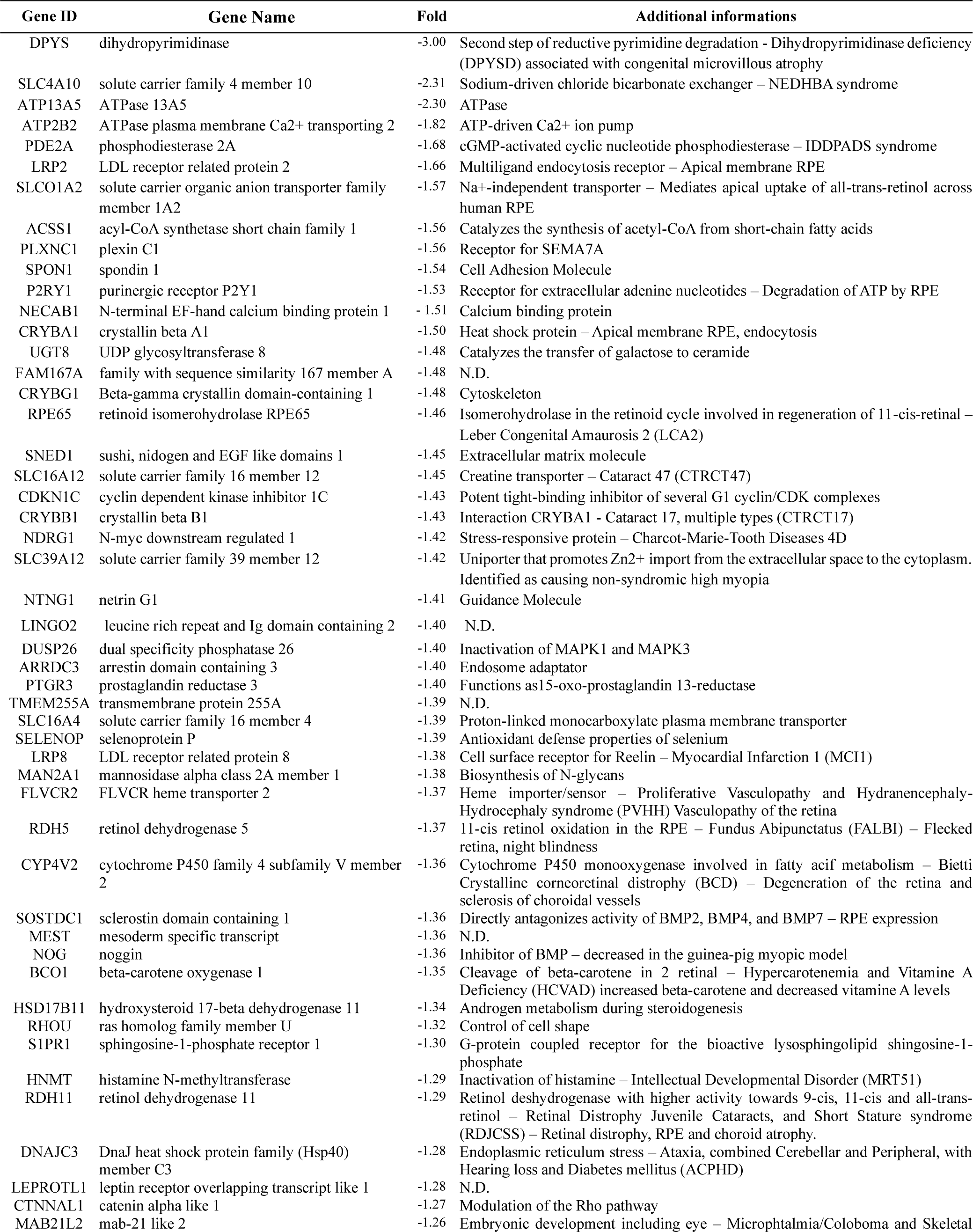
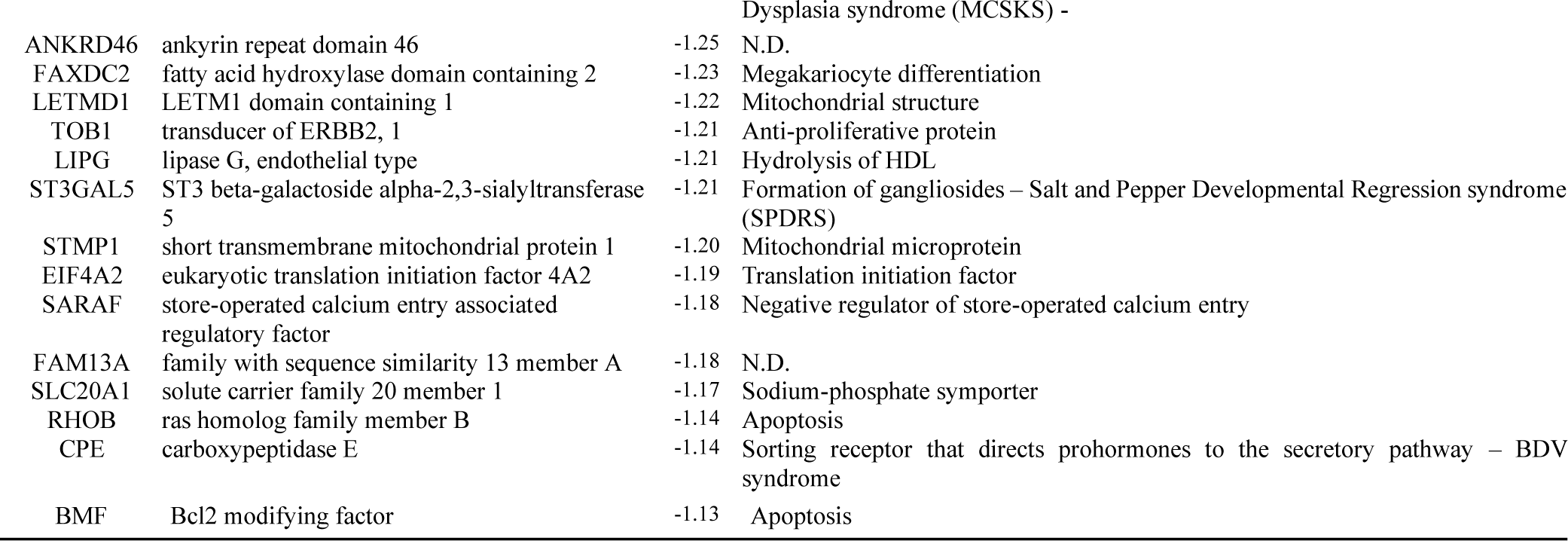
List of differentially expressed genes (DEGs) down regulated in shLRP2 iRPE cells.

**Table 4.**
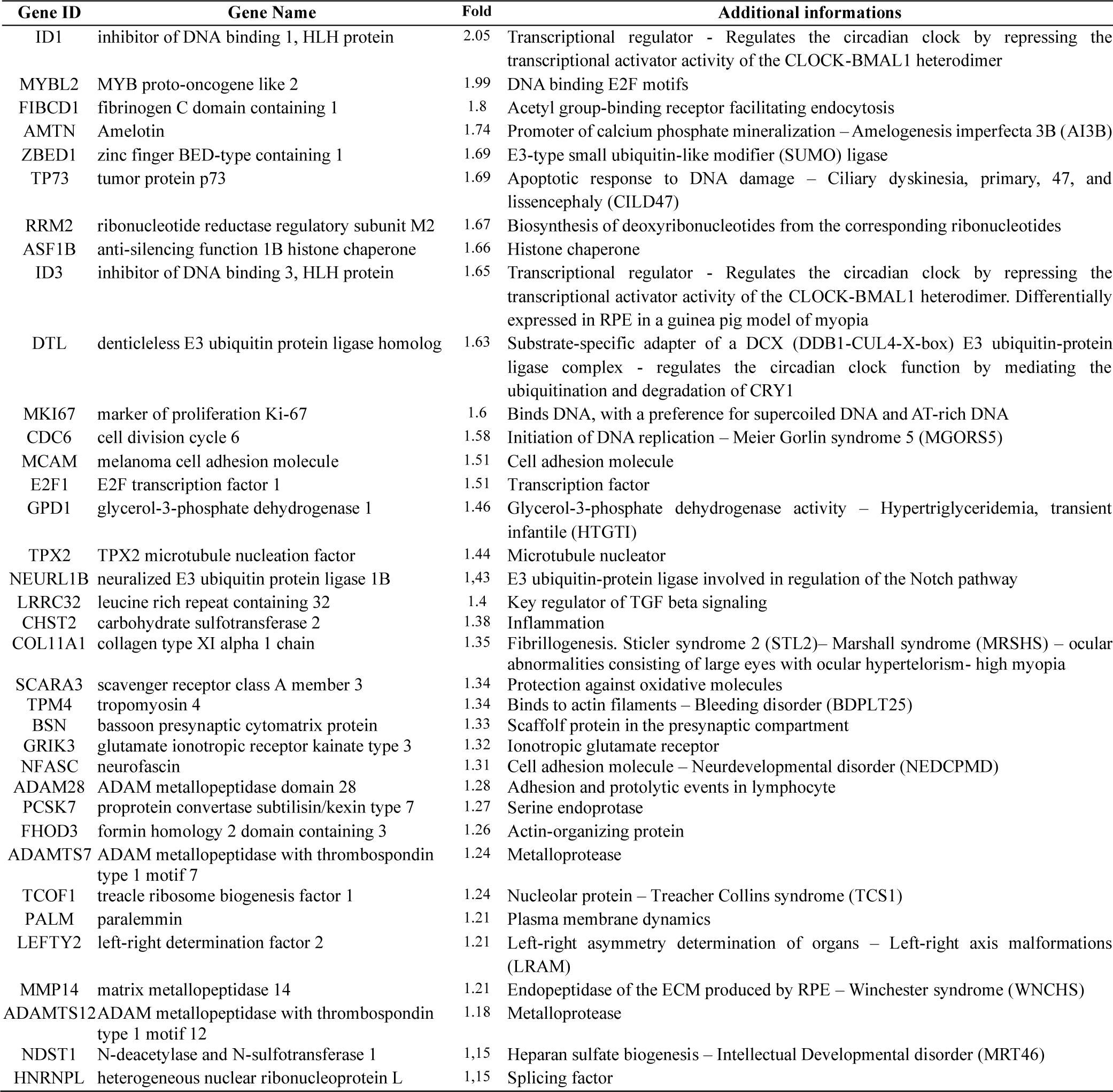
List of DEGs up regulated in *in shLRP2* iRPE cells with a fold change cut off of 1.1<.

**Table 5.** Selected gene sets associated with LRP2 reduction in iRPE cells. Normalized Enrichment Scores.

| ID | Description | NES |
| --- | --- | --- |
| **Light processing** |  |  |
| M11434 | GOBP RETINOL METABOLIC PROCESS | -2,291 |
| M13006 | GOBP SENSORY PERCEPTION OF LIGHT STIMULUS | -2,269 |
| M34809 | HP SCOTOMA | -1,966 |
| M38378 | HP ABNORMAL FULL FIELD ELECTRORETINOGRAM | -1,952 |
| M37110 | HP PROGRESSIVE NIGHT BLINDNESS | -1,919 |
| M14699 | GOBP SENSORY PERCEPTION | -1,910 |
| M41363 | HP ABNORMAL LIGHT AND DARK ADAPTED ELECTRORETINOGRAM | -1,893 |
| M37157 | HP PERIPHERAL VISUAL FIELD LOSS | -1,855 |
| **Hormonal processes** |  |  |
| M12960 | GOBP RESPONSE TO STEROL | -2,220 |
| M12703 | GOBP CELLULAR HORMONE METABOLIC PROCESS | -2,075 |
| M14197 | GOBP CELLULAR RESPONSE TO STEROL | -2,032 |
| M3753 | GOBP HORMONE METABOLIC PROCESS | -1,873 |
| M11178 | GOBP REGULATION OF HORMONE LEVELS | -1,681 |
| M16758 | GOBP STEROID METABOLIC PROCESS | -1,631 |
| M13381 | GOBP STEROID BIOSYNTHETIC PROCESS | -1,592 |
| **Epithelial transport** |  |  |
| M37121 | HP RETINAL PIGMENT EPITHELIAL ATROPHY | -2,029 |
| M13303 | GOBP TRANSEPITHELIAL TRANSPORT | -2,017 |
| M24808 | GOBP IMPORT ACROSS PLASMA MEMBRANE | -1,921 |
| M15727 | GOBP IMPORT INTO CELL | -1,748 |
| M18079 | GOMF METAL ION TRANSMEMBRANE TRANSPORTER ACTIVITY | -1,702 |
| M10755 | GOBP TRANSITION METAL ION TRANSPORT | -1,668 |
| **Apico-basal polarity** |  |  |
| M1446 | GOCC APICAL PLASMA MEMBRANE | -1,699 |
| M17164 | GOCC APICAL PART OF CELL | -1,698 |
| M3406 | GOCC BASOLATERAL PLASMA MEMBRANE | -1,689 |
| M17798 | GOCC BASAL PART OF CELL | -1,686 |
| **Choroid support** |  |  |
| M34087 | GOBP VASCULAR TRANSPORT | -2,088 |
| M34782 | HP CHORIORETINAL ATROPHY | -2,039 |
| M38919 | HP CHORIORETINAL DEGENERATION | -1,904 |
| M16767 | GOBP VASCULAR PROCESS IN CIRCULATORY SYSTEM | -1,843 |
| **Extracellular matrix remodeling** |  |  |
| M12927 | GOMF EXTRACELLULAR MATRIX STRUCTURAL CONSTITUENT | 2,155 |
| M10742 | GOCC COLLAGEN TRIMER | 2,154 |
| M25882 | GOCC COLLAGEN CONTAINING EXTRACELLULAR MATRIX | 2,060 |
| M10505 | GOBP COLLAGEN FIBRIL ORGANIZATION | 1,995 |
| M23167 | GOBP COLLAGEN METABOLIC PROCESS | 1,983 |
| M17310 | GOCC COMPLEX OF COLLAGEN TRIMERS | 1,949 |
| M22974 | GOBP COLLAGEN CATABOLIC PROCESS | 1,869 |
| M880 | GOBP PROTEOGLYCAN METABOLIC PROCESS | 1,835 |
| **Cytoskeleton assembly** |  |  |
| M6458 | GOCC ACTIN FILAMENT | 2,351 |
| M17227 | GOCC ACTIN FILAMENT BUNDLE | 2,125 |
| M17139 | GOCC MYOFILAMENT | 2,059 |
| M17284 | GOCC ACTOMYOSIN | 2,032 |
| M5272 | GOCC ACTIN CYTOSKELETON | 2,011 |
| M18964 | GOMF MICROTUBULE MOTOR ACTIVITY | 2,010 |
| M9766 | GOMF ACTIN FILAMENT BINDING | 1,915 |
| M14799 | GOMF ACTIN BINDING | 1,895 |
| M9078 | GOMF MICROTUBULE BINDING | 1,888 |
| M2926 | GOCC CORTICAL CYTOSKELETON | 1,865 |
| M4679 | GOCC MICROTUBULE ASSOCIATED COMPLEX | 1,864 |
| M12616 | GOBP MICROTUBULE DEPOLYMERIZATION | 1,862 |
| M14007 | GOBP MICROTUBULE BUNDLE FORMATION | 1,830 |

### Light and cortisol regulate *LRP2* expression in human iRPE cells

Light and circadian rhythms are major environmental factors associated with the incidence and the progression of myopia^26^ and cortisol secretion which is under circadian regulation is modulated by light exposure^27^. We assessed the exposure of iRPE cells to an LED lighting system emitting red [631nm], blue [454nm], and hot white [3300°K] light at a dose of 0.3 J/cm², which was previously shown to be safe for the iRPE^28^. After 0.5 and 2 hours of exposure to any of these LEDs, the expression of LRP2 mRNA increased significantly (2.00 ± 0.16, *p* = 0.0059 and 1.70 ± 0.08, *p* = 0.0132) and transiently, returning to baseline at 10 hours (1.02 ± 0.18, *p* = 0.9791) (**Fig. 8A**). Exposure to red light was the most effective at increasing LRP2 mRNA at 0.5 and 2 hours (2.08 ± 0.14, *p* = 0.0286 and 1.83 ± 0.06, *p* = 0.0286) (**Fig. 8A**). On immunohistochemistry, two hours after illumination, we observed a parallel increase in LRP2 and the early endosome-associated protein 1 (EEA1) in iRPE cells (**Fig. 8B**).

**Fig. 8.**
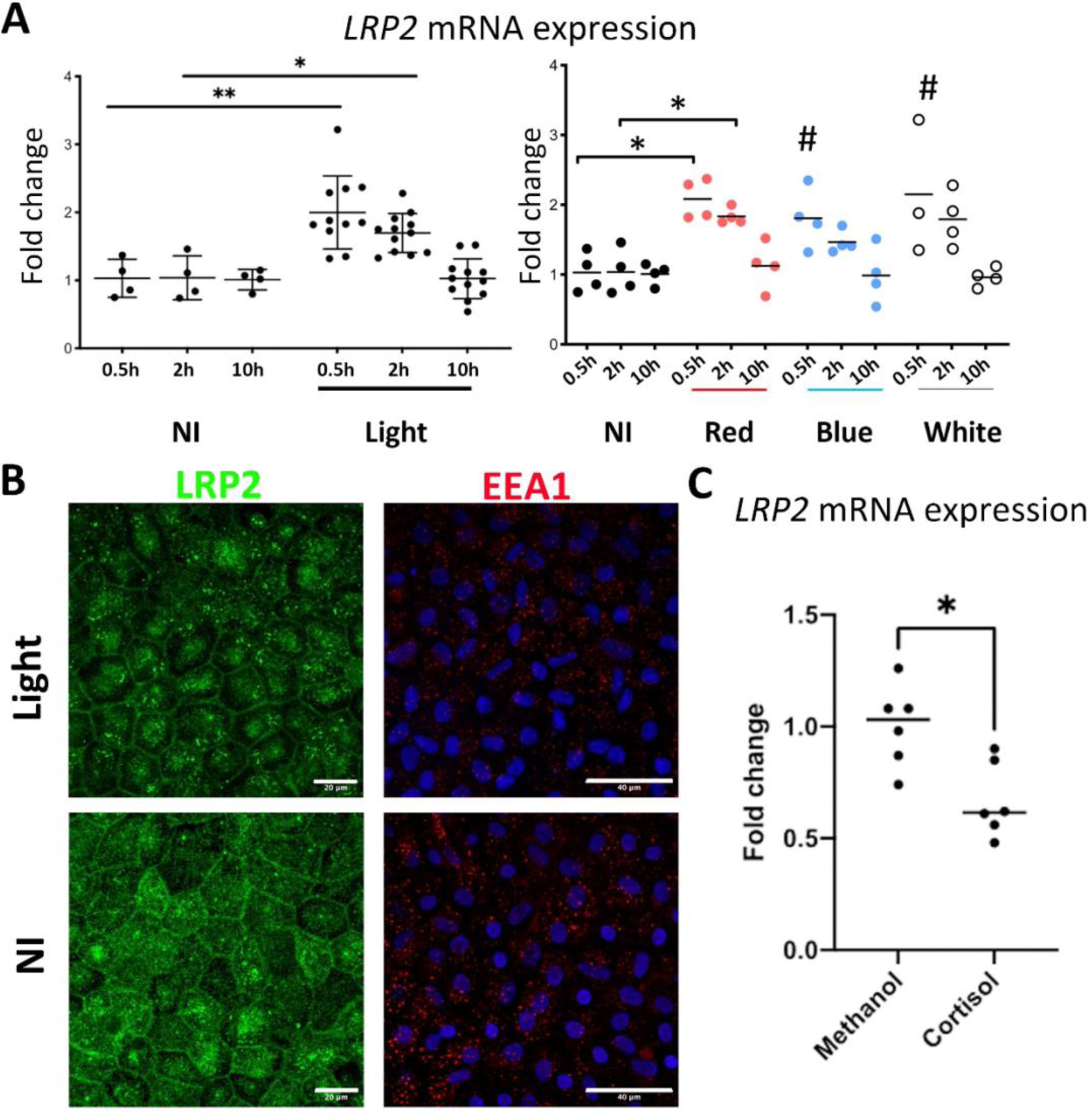
**LRP2 expression and environmental factors**. (**A**) Quantification of *LRP2* mRNA by Q-PCR in iRPE non-exposed (ne) or exposed to light (left panel) after 0.5, 2 or 10 hrs post illumination. Left panel showing quantification of *LRP2* mRNA in iRPE non-exposed (ne) or exposed to light. Right panel showing quantification of *LRP2* mRNA in iRPE non-exposed (ne) or exposed to red, blue, or white light. Values correspond to the mean of 4 independent experiments in duplicates for each condition. Each independent experiment represents the mean of three wells. Datas were expressed in fold gene expression ± SD. Data were analyzed using the non-parametric Kruskal Wallis test and Mann Witney post-test, \**p*<0.05; \*\**p*<0.01, # not significant. (**B**) LRP2 and EEA1 expressions in iRPE non-exposed or exposed to red light. (**C**) *LRP2* mRNA expression in iRPE cultures treated without (ethanol) or cortisol. Values correspond to the mean of 6 experiments in triplicates. Datas represent mean fold gene expression ± SEM. Mann-Whitney test, \**p*<0.05. Scale bars: 20 μm for LRP2 and 40 μm for EAA1.

Having previously demonstrated that the glucocorticoid receptors NR3C1 (GR) and the mineralocorticoid receptor NR3C2 (MR) are both expressed and functional in the iRPE cells used in this study^29^, we evaluated the effect of cortisol on *LRP2* expression. Exposure of iRPE cells to 100nM cortisol led to a significant decrease in LRP2 mRNA expression after 24 hours (1.00 ± 0.04 vs. 0.67 ± 0.07; p = 0.0152*<0.05) (**Fig. 8C**).

Both light exposure and cortisol regulate the expression of *LRP2* in iRPE cells but in opposite manner, suggesting that LRP2 could be one of the molecular links between environmental factors and HM development.

## Discussion

Convergent results from various experimental models have previously identified LRP2 as an important player in eye growth and in staphyloma. Both the *Lrp2*-cKO mouse and the conditional mouse with specific knock out of *Lrp2* in RPE developed myopia and staphyloma^15,16^. Of note, the latter models are the only myopia models that develop staphyloma, showing the important role of LRP2 expressed in the RPE in its development. In humans, biallelic pathogenic variants in *LRP2* cause the Donnai-Barrow syndrome characterized among others signs by high grade syndromic myopia and PS^14^. In these patients LRP2 is either absent or nonfunctional^30,31^ indicating that LRP2 is important in human eye pathophysiology. We here show that reduced levels of LRP2 in the vitreous, retina and RPE cells are associated with NSHM, supporting the hypothesis that LRP2 is also involved in NSHM associated with PS in humans. The exact roles of LRP2 in human retina physiology and its mechanisms of action on eye growth are not fully elucidated. Nevertheless, it is likely that any changes in LRP2 synthesis, recycling and/or subcellular distribution could alter its functions. Indeed, studies in the renal proximal convoluted tubule, the major site of LRP2 expression in the adult, show that in its membrane bound form LRP2 acts as major endocytic receptor whereas processed by regulated intramembrane cleavage, the LRP2 C-terminal fragment would enter the nucleus to regulate gene expression^32^. Extracellular cleaved LRP2 fragment is thought to act as a soluble decoy receptor that may bind ligands and alter their bioavailability ^18,33^.

The present data suggest that both the membrane bound and the extracellular form of LRP2 are decreased in the myopic eyes. The reduction of soluble LRP2 in the vitreous, shown by two different biochemical methods, performed by two independent groups in two different populations together with reduced LRP2 observed in the retina and RPE suggests that the reduction of LRP2 results from a decrease in transcription and / or translation. Although soluble LRP2 has been previously measured in human vitreous proteome^34^ its origin is unknown. In renal cells, a soluble form of LRP2 has been detected, resulting either from the secretion of an isoform lacking the cytoplasmic domain or from a two steps cleavage by γ-secretase for intramembrane cleavage and by metalloproteases for the extracellular cleavage ^35^. The detection of LRP2 in the vitreous and the plasma membrane/endocytic apparatus of retinal cells in both control and myopic subjects may suggest that extracellular LRP2 cleavage commonly occurs in the ocular tissues. Interestingly, transthyretin, that indirectly influences the cleavage of LRP2 by γ- secretase by modulating its interaction with the receptor, was shown to be misfolded and dyfunctional in case of myopic macular pathologies^36^. The exact functions of the soluble form of LRP2 still remain to be understood. It could act as a ligand trap, masking or enhancing signaling pathway such as that of SHH^37^ or BMP4^22^ that was associated with HM genetic studies^38^. Indeed, excessive levels of soluble LRP2 induced by a CRISPR/Cas9 approach were associated with buphthalmia in the zebrafish^39^. The authors suggested that defective ligand endocytosis resulting from a lack or excess of LRP2 resulted in the same eye enlargement, but further studies are necessary to identify the underlying mechanism.

At the cellular level, LRP2 was expressed mostly in the apical and basal extremities of glial Müller cells in the human retina. Recently, LRP2 was shown to be expressed in brain atsrocytes and microglia, differentially regulated by inflammatory signals and involved in amyloid beta endocytosis activity^40^. In the RPE, LRP2 was located at the apical membrane and in vesicles following a decreasing gradient along the apico-basal axis, ensuring transport between the subretinal space and the choroid as previously suggested ^17^. We also identified LRP2 signal at the basal side underneath RPE in the normal human retina, where it could bind to clusterin and apolipoproteins and metalloproteases, abundant in the Bruch membrane in humans^41^. LRP2 expressed at the baso-lateral could contribute to the regulation of nutrient diffusion and waste disposal between the RPE and blood ^41^, eye homeostasis and eye growth ^17,42^. In NSHM RPE LRP2 expression was reduced along the endocytic apparatus suggesting defective LRP2 - dependent endocytosis. LRP2 is considered as a suppressor of the eye size; the exact mechanism is not clear but reduced endocytosis and recycling of LRP2 and LRP2-ligand complexes is likely to cause a less-efficient LRP2-mediated regulation of eye growth. Reduced LRP2-ligand endocytosis by RPE cells would eventually result in an excess of these ligands in the milieu. We did not observe any upregulation of transthyretin, LRP2 ligand previously involved in HM^36,43^, although the levels of clusterin were slightly increased. On the contrary RBP4, another LRP2 ligand was upregulated (>1.5 fold) suggesting a dynamic regulation of these proteins in the vitreous.

To further investigate the relationship between myopia, SP, and LRP2 expression in RPE cells, we analyzed the transcriptomic changes induced by partial knockdown of LRP2. Our study also shows that LRP2 depletion in the RPE may lead to a coordinated modulation of sensory perception and phototransduction with that of hormone and steroid metabolism and with tissue remodeling. The decrease in transcripts of enzymes related to retinoid homeostasis (RPE65, retinal dehydrogenase 5 or 11) may explain the decrease in visual cycle integrity and the retinal degeneration, observed in humans and in animal models null for *Lrp2*^44^. In addition to the differentially apically expressed solute transporters, we found numerous genes involved in maintaining the epithelial apical scaffold, such as occluding (OCLN) or myosin VIIA (MYO7A). Both in the Foxg1-Cre-Lrp2lox/lox mouse eye and in human eyes with PS, RPE cell structure and polarity were severely altered, which could be a consequence of ocular elongation, but also an early and possibly LRP2-driven event, as shown in form-deprived myopia in chicks. In the latter, an increase in the surface area of individual RPE cells compensated for the expanded vitreous chamber^45^. Similarly, in the Foxg1-Cre-Lrp2lox/lox mouse, we showed in previous studies that the total surface area of the RPE layer was expanded without RPE proliferation or cell death^15,46^. We observed that LRP2 deficiency leads to a severe reduction in clathrin-mediated endocytosis vesicles in humans and in iRPE cells, together with a reduction in EEA1 and LAMP1 vesicles. We could thus speculate that the loss of LRP2 in RPE cells could induce cell surface enlargement through failure of apical membrane removal. As a result of surface expansion, especially posterior to the equator, the RPE would then produce more Bruch’s membrane, which is consistent with morphometric changes of Bruch’s membrane in the development of myopia in humans^47^. However, further studies are required to better understand the mechanisms linking LRP2 expression in RPE cells and Müller glial cells to the development of myopia and PS.

The causes of LRP2 downregulation in myopic eyes is most probably multifactorial and could result from tissue scale mechanical stresses^48^ occurring during excessive eye growth, from genetic or epigenetic predispositions, from local inflammatory microenvironment^49,50^, from hormonal factors and, from environmental stimuli such as lighting conditions. *LRP2* expression is tightly regulated and its reduction in epithelial cells has been associated in different fibrosis pathologies. Further, this inhibition of transcription is caused by the canonical TGF-beta1/SMAD2-SMAD3 pathway^51^. Among the factors that positively control *LRP2* are retinoic acid^52^, vitamin D^52^, and alpha and gamma peroxisome proliferator-activated receptors^53^. Interestingly, our results show that *LRP2* expression is upregulated in iRPE by light exposure, which can be explained by the presence of melanopsin (Opn4) ^54,55^ and other non-visual opsins like neuropsin (Opn5) ^56^ or peropsin ^57^ in RPE cells. *LRP2* up-regulation by light, is in line with epidemiological data showing that light exposure is a robust environmental stimulus in the prevention of myopia^58^. Our experiments indicated that exposure to red light seemed to up-regulate *LRP2* more efficiently than blue or white light, in line with intervention studies in children showing reduced myopia progression by red light therapy ^59^. An indirect mechanism of action of light could be the circadian regulation of cortisol that is sensitive to the wavelength composition of environmental lighting^27^. Interestingly, cortisol downregulated the expression of LRP2 in iRPE cells and in a Chinese population, higher levels of cortisone, corticosterone and aldosterone were recently measured in aqueous humor of myopic eyes^60^, suggesting a link between myopia and ocular metabolism of corticoids. In a model of negative lens induced myopia in guinea pigs, hydrocortisone enhanced the axial elongation, the myopic shift and scleral thinning but had the reverse effect during physiologic emetropization^61^, demonstrating the potential differential effects of corticoids in emmetropization process and in pathologic myopia. The complex links between light, circadian rhythm, corticoids, LRP2 myopia require further investigations but our results tend to indicate that LRP2 could be a molecular link between light, circadian regulations and NSHM.

This study is subject to certain limitations, including the relatively small number of human eyes with PS that were available for analysis due to the rarity of such fresh mortem eyes in biobanks. However, the observation that LRP2 was not only reduced in the retina but also in the vitreous lends further support to our findings. The number of vitreous samples is also limited; however, this study has examined a substantial number of vitreous samples from HM patients with PS, and the findings have been validated by the reproducibility of the results by two independent groups in two distinct patient populations. Subsequent studies should aim to specifically analyze the relationship between LRP2 in the vitreous, axial length, and PS, and to characterize the soluble LRP2 form.

In conclusion, our study demonstrates that in human eyes with NSHM associated with PS, which is the more severe form of myopia, LRP2 is decreased both in the vitreous and in the RPE. In human RPE cells, LRP2 expression is regulated by light, which is the environmental factor the most associated with myopia. Finally, the silencing of *LRP2* in human RPE cells regulated the expression genes involved in myopia development. LRP2 appears as a potential interventional target in the prevention of NSHM and its blinding complications.

## Materials and methods

### Ethics - Patients recruitment

A total of 25 patients with NSHM and 23 patients with emmetropic eyes, for which vitrectomy was scheduled for macular surgery or for intraocular lens luxation were recruited in the study. There were 25 Asian Japanese patients and 23 Caucasian European patients. Undiluted vitreous humor samples from eyes with NSHM complicated by staphyloma (n=15) and from emmetropic control eyes (n=10) were obtained for mass spectrometry analysis at Kyoto Prefectural University of Medicine, Japan (**Table 1**). The study was conducted in compliance with the Institutional Review Board of Kyoto Prefectural University of Medicine which approved the study (permission RBMR-C-864-6). Informed written consent was obtained from all patients. Undiluted vitreous humor samples from eyes with NSHM and PS (n=10) and from emmetropic control eyes (n=10) were collected for ELISA analysis at Ophtalmopole, Paris, France and at Hospital Puerta de Hierro, Albacete, Spain (**Table 1**). In France and Spain, vitreous samples are considered as surgical waste and can be used for research purposes if patients have expressed no opposition according to French and Spanish law. Agreement was obtained for all patients. Storage and analysis were. authorized in France by CPP Ile de France 1 (N°2016-14390) and the use of patient’s information got the authorization number CNIL 2233436. All samples were stored at -80°C until preparation was initiated.

### Sample Preparation for Mass Spectrometry

Protein concentrations were measured on undiluted vitreous samples with an infrared spectrometer (Direct Detect, Darmstadt, Germany) and samples were prepared for analysis using the S-Trap Micro spin column digestion protocol from ProtiFi (Huntington, NY, USA) as described in detail^62^ . Analysis was performed on an Orbitrap Fusion Tribrid mass spectrometer coupled to a Dionex UltiMate 3000 RSLC nano system (Thermo Fisher Scientific Instruments, Waltham, MA, USA). All details are described in the **Supplemental section of materials and methods**.

### Enzyme-Linked Immunosorbent Assay (ELISA) protein quantification of LRP2

LRP2 concentration was determined by ELISA (Creative Diagnostics, DEIA-FN834, NY). Briefly, microtiter plates were coated with 1μg polyclonal antibody specific for human LRP2. Undiluted 100 µl of aqueous or vitreous humor sample was added to the plates. The presence of LRP2 was revealed by incubation with the same antibody labeled with biotin. The intensity of this colored product is directly proportional to the concentration of LRP2 present in the samples and was measured immediately by absorbance at 450 nm. The Student’s *t*-test was used for statistical analysis of the ELISA data. Differences and correlations were considered statistically significant if *P* <0.05. Two-way scatter plots based on ELISA data with prediction from a linear regression were created in STATA 16.0.

### Eye samples

Four donor eyes were obtained by Lion’s Gift of Sight (Saint Paul, Minnesota, USA), which operates under the rules of the Food and Drug US Administration and the Eye Bank Association of America. Donor consent was obtained, and the next of kin received no compensation or financial gain from the donation. Samples were collected less than 10 hours after death and fixed in 4% paraformaldehyde (PFA) in PBS for 24 hours. Importation into France was carried out in accordance with regulations applicable to the transfer of human tissues with authorization N° IE-2022-2285 from the French ministry of research. Two male donor eyes with moderate cataract and refraction of -0.75 (+0.75)5° in the right eye and -0.50 (+1) 170° in the left eye were used as emmetropic control eye. The donor aged 88 male no remarkable medical history and no known ocular diseases and died due to small bowel obstruction. The postmortem time before enucleation and fixation was 6 hours.

Both HM eyes from a 89 male donor were analyzed. Medical history reported hypertension and arthritis, the cause of death was pulmonary edema and postmortem time was 8 hours. Both eyes were pseudophakic and showed macular staphyloma, vitreomacular traction and degenerative macular schisis. Pre mortem examination of the left eye was impaired by corneal opacification secondary to corneal surgery for keratoconus. Eyes were fixed in 4% paraformaldehyde overnight, rinsed and preserved in 1% paraformaldehyde at 4°c. Eyes were then dissected to remove the cornea and the anterior segment. The posterior segment was included in Optimal cutting temperature compound and 10µm cryosections were prepared. For flat-mounting, after removing the cornea and lens, the remaining posterior segment of the eye was flat-mounted and dissected to remove the neural retina. The posterior segment was cut into 4 quarters and the RPE/choroid was separated from the sclera after section of the vortex veins.

### Immunostaining of human tissues

Sclera-choroidal-RPE complexes were incubated for 1 hour in a solution (phosphate-buffered saline (PBS) 0.1M, 10% Normal goat serum, 0.01% triton X100) at room temperature. Then, tissues were incubated with the primary antibodies at appropriate dilution (Table 6) and phalloidin-rhodamine (1:200, Thermo Fisher Scientific, France) in a buffer solution (PBS 0.1M, 5% Normal donkey serum, 0.01% triton X100) during 5 days at 4°C under gentle agitation. After 6 X 30 min consecutive washings with 0.1M PBS / 0.01% triton X100, tissues were incubated for 3 hours at room temperature with the adequate secondary antibodies diluted at 1:1000. After 4 X 30min successive washings, tissues were flat mounted using Dako Omnis Fluorescence Mounting Medium (Agilent, Les Ulis, France). Similar protocol was applied for immunostaining on transversal cryosections, with overnight (4°C) incubation for the primary antibody and 1 hour (at room temperature) for secondary antibody incubation. Images were acquired using a fluorescence microscope (model Olympus BX51, Olympus, Rungis, France) or a confocal microscope (model Zeiss LSM710, Zeiss, Rueil Malmaison, France).

**Table 6.** . Antibodies and reagents for immunohistochemical staining.

| Primary Antibodies | Species, Label, Method | Source, number | Dilution |
| --- | --- | --- | --- |
| Glutamine synthetase | Mouse, IF | Merck Millipore, MAB2-302 | 1:300 |
| Tubulin-beta3 | Mouse, IF | BioLegend, 801202 | 1:300 |
| OPN1LW/OPN1MW (Red/green-sensitive opsin) | Rabbit, IF | Merck Millipore, AB5405 | 1:100 |
| OPN1SW (Blue-sensitive opsin) | Goat, IF | Santa Cruz, sc-14363 | 1:200 |
| Glial fibrillary acidic protein | Rabbit, IF | Dako Cytomation, Z0334 | 1:300 |
| LRP2 | Rabbit, IF | Abcam, Ab76969 | 1:300 |
| ZO-1 | Rabbit, IF | Invitrogen, 40-2200 | 1:200 |
| Clathrin | Mouse, IF | Santa Cruz, sc-12735 | 1:200 |
| EEA1 | Mouse, IF | BD Transduction, 610457 | 1:250 |
| LAMP1 | Mouse, IF | DSHB, H4A3 | 1:100 |
| **Secondary Antibodies** |  |  |  |
| Anti-Rabbit | Donkey, alexa-488, IF | ThermoFisher, A21206 | 1:300 |
| Anti-Rabbit | Donkey, alexa-594, IF | ThermoFisher, A21207 | 1:300 |
| Anti-Mouse | Donkey, alexa-594, IF | ThermoFisher, A32766 | 1:300 |
| Anti-Mouse | Goat, alexa-594, IF | ThermoFisher, A11005 | 1:300 |
| **Reagents** |  |  |  |
| Phalloidin (Aminata Phalloidin) | Rhodamine, IF | ThermoFisher, R415 | 1:300 |
| 4',6-diamidino-2-phenylindole (DAPI) | Fluorescent counterstain | Roche, 10236276001 | 1:2500 |

### iRPE cell culture and differentiation

A human-induced pluripotent stem cells (hiPSC) cell line, obtained from fibroblasts of a healthy donor was used as previously described ^29^. The hiPSC were expanded and differentiated into hiPSC-derived RPE (iRPE) cells using a differentiation protocol^29^. All details of cell culture and maintenance are described in the **Supplemental section of materials and methods**.

At passage 3, iRPE cells were seeded in Transwells (0.4 μm pore polyester membrane inserts, Corning, CLS3450 and CLS3460) coated with Matrigel® red phenol-free matrix (Corning, 356231) in a serum- and antibiotic-free retinal Retinal Differentiation Medium (**Supplemental section of materials and methods**). All cultures were maintained in an incubator at 37 °C, 5% CO2. The trans-epithelial resistance (TER) was measured (**Supplemental section of materials and methods**), and cells were used between the sixth and the tenth week of culture, once the trans-epithelial resistance reached physiological levels (> 100 Ω/cm^2^).

### RNA silencing

Briefly, for each transfection, 6µl of triplex siRNA, a pool of 3 target-specific 19-25 nucleotides *siLRP2* (Santa Cruz, sc-40103) or scrambled *siRNA* (Santa Cruz, sc-37007) or fluorescein-conjugated control siRNA (Santa Cruz, sc-36869) were diluted in 100µl of siRNA transfection medium (Santa Cruz, sc-36868,). This solution was mixed v/v with a diluted solution of siRNA transfection reagent (Santa Cruz, sc-29528). The siRNA mixture was added to the apical surface of Transwell-coated iRPEs for 6 hours. Without removing the siRNA mixture, serum- concentrated medium (2X) was added for 18 h incubation. After incubation, the cells were placed in fresh medium for 24 hours. The cells were then collected for protein and RNA extraction.

### RNA sequencing

Raw data quality was assessed using FastQC. Low-quality sequences and adapters were pruned or removed using Dragen with default settings to retain high-quality matched reads. Illumina’s DRAGEN Bio-IT platform (v3.10.4) was used for mapping to the hg38 reference genome and quantification using the Gencode v37 annotation GTF file. Library orientation, composition and transcript coverage were checked using Picard tools. Subsequent analyses were performed using R software. Data normalization was performed using the DESeq2 (v1.26.0) Bioconductor software package, followed by differential expression analysis using the DESeq2 workflow. Adjusted p-values for multiple hypotheses were calculated using the Benjamini-Hochberg procedure to control the false discovery rate (FDR). Finally, an enrichment analysis was performed using the R clusterProfiler (v3.14.3) package for Gene Set Enrichment Analysis (GSEA) on gene sets from the C5 ontology, comprising GO (Gene Ontology) grouping biological processes, cellular components and molecular functions and HPO (Human Phenotype Ontology), and the KEGG and REACTOME databases.

### Light exposure

Cells were exposed to red (630 nm, Qasim1xQA-SL0001-EU), blue (454 nm, Joyland, D50SWD-B), or white (3300 K, Lepro4100067-WW-EU-NF-a) LEDs in a 37°C incubator with 5% CO2. The specificities of each light source (CT, peak of emission, integrated irradiance) were measured using a spectroradiometer (Kanonica Minolta CL70 F CR I). Cells were exposed to a white, blue and red LED for 820s, 1000s and 900s respectively, to match ambient light exposure. These exposure times correspond to a dose of 0.3J/cm2 (calculated as the energy of the integrated spectrum multiplied by the exposure time). Cells were illuminated for 5 days, and mRNA collected at 3 post-illumination times: 30 minutes, 2 hours and 10 hours. TER measurements were taken during the blue-light illumination protocol to assess cell integrity during the experiment.

### Corticosteroid treatment of iRPE cells

Cells were seeded at P3 in cell culture plastic dishes. On day 35, one week prior to corticosteroid treatment, retinal differentiation medium was removed and iRPE cells were incubated in experimental corticosteroid-free medium (DMEM, high glucose, HEPES, no phenol red (Thermo Fisher Scientific,); 10% fetal bovine serum, charcoal stripped (Thermo Fisher Scientific)). On day 42, iRPE cells were treated for 24 h with cortisol 10^−7^ M. As cortisol was dissolved in ethanol (EtOH), control cells were treated with 0.1% EtOH in medium **Quantitative RT-qPCR**

Total RNA was isolated using commercially available kits according to the manufacturer’s instructions (RNeasy Mini, Qiagen) and measured (Nanodrop, Peqlab). One μg was used in a reverse transcription reaction (SuperScript First strand synthesis, ThermoFisher). Quantitative PCR was performed using Master Mix PCR Power SYBR™ Green (4367659, Applied Biosystems). Quantitative polymerase chain reaction (PCR) was performed on a 7500 real-time PCR system (Applied Biosystems). Transcripts levels were calculated using the standard curves generated using serial dilutions of cDNA obtained from samples, then normalized to HPRT. Primers sequences were listed in Table 6. Each plotted value corresponds to the mean of 3 independent experiments in duplicates Datas represent a mean gene expression per fold ± s.d. Data were analyzed using the non-parametric Kruskal Wallis test and Mann Witney post-test (*p<0.05, **p<0.01, ***p<0.001), using GraphPad Prism 8 software (GraphPad Software, San Diego, CA).

### Immunocytochemistry on iRPE cells

iRPE cells were fixed with 4% PFA. Primary antibodies were incubated in blocking buffer (1% BSA, 0.1% triton X-100 in PBS) for three hours. Secondary antibodies were incubated for one hour. Nuclei were counterstained with DAPI for 5 minutes. Antibodies and reagents listed in **Table 6**. No cellular autofluorescence or non-specific labeling was detected under these conditions. Images were collected by confocal microscopy and processed using ZEN (Zeiss) and ImageJ softwares.

### Immunofluorescence

Eyes were enucleated (n = 7 WT, n = 7 Lrp2-cKO; age = 20 weeks) and fixed for 30 minutes in 4% PFA. The RPE was dissected and laid flat, then post-fixed with acetone for 10 minutes at -20°C.. Tissues were incubated with phalloidin for 3 h, rinsed, and then incubated with rabbit anti-ZO1in blocking buffer (1% BSA, 0.3% Triton X-100 in PBS) for 24 h at 4°C. Secondary antibody was incubated 2 h at room temperature. After mounting, RPE was cut at four spots yielding four quadrants corresponding to the naso-temporal and dorso-ventral directions. Images were acquired using an inverted confocal microscope (Zeiss LSM 710). Flatmount was imaged with a 20x objective lens. Typically, 3 to 4 images were obtained of each of the four quadrants to cover studied regions (intermediate or peripheral retina). Using ImageJ software, a macro was developed to calculate the RPE cell areas for each sample. . The macro is divided into three main stages: the first stage involves segmentation, including image binarization and skeletonization; the second stage involves manual ROI correction; and the final stage involves skeleton analysis, requiring the following plugins: tubeness, morphology, and analyze skeleton. To calculate the RPE cell density, the total area of RPE cells cells per region of interest (intermediate or periphery) was divided by the total number of RPE cells per region of interest. Data were analyzed using the non-parametric Mann Witney post-test using GraphPad Prism 8 software (GraphPad Software, San Diego, CA).

## List of supplementary materials

### Supplemental Material and methods

#### 5 Supplemental Figures

Figure S1: GO analysis of DEPs

Figure S2: The most significant cluster obtained from the PPI and its disease enrichment Figure S3: Principal component analysis.

Figure S4: KEEG, Gene Ontology, Reactome and Human Phenotype Ontology based gene enrichments.

Figure S5: Category net plot enrichment analysis.

#### 4 Supplemental Data Files in attached Excel Format files

Data File S1: filtered results of the LC/MS MS

Data File S2: List of DEGs

Data File S3: Enrichment-*p* adjust

Data File S4: Enrichment LRP2-*p* adjust

## Author Contributions

KD, AT, OC and FBC: study conception and design - writing original draft. KD, E Picard, PL, LJ and E Pussard: performing experiments. RK: providing Lrp2-cKo. JMRM and JRM: providing vitreous samples. KK collected vitreous samples for mass spectrometry and contributed to the analysis of proteomics data. HV, BH, and LJC performed mass spectrometry and data analysis of the proteomics data. All authors contributed to review and editing of the article and approved the submitted version.

## Institutional review Board Statement

At Kyoto Prefectural University of Medicine, Japan, the study was conducted in compliance with the Institutional Review Board of Kyoto Prefectural University of Medicine which approved the study (permission RBMR-C-864-6). Informed written consent was obtained from all patients. At Ophtalmopole, Paris, France, vitreous samples are considered as surgical waste and can be used for research purposes if patients have expressed no opposition according to French law. Agreement was obtained for all patients. Storage and analysis were authorized in France by CPP Ile de France 1 (N°2016-14390) and the use of patient’s information got the authorization number CNIL 2233436.

At Hospital Puerta de Hierro, Albacete, Spain, vitreous samples are considered as surgical waste and can be used for research purposes if patients have expressed no opposition according to Spanish law. The study involving human iRPE cultures were reviewed and approved by CODECOH DC- INSERM. Importation of human eyes was approved by the French ministry of research (N° IE-2022-2285)

## Informed Consent Statement

Informed consent to participate in this study was obtained from all the participants.

## Conflict of interest

The authors declare that the research was conducted in the absence of any commercial or financial relationships that could be construed as a potential conflict of interest.

### Acknowledgments

We gratefully acknowledge the support of the Data Analysis Core (DAC) platform at the Institut du Cerveau de Paris in carrying out RNA sequencing. Special thanks also go to Emeline Cherchame for her invaluable help with RNAseq data analysis. The services provided by the DAC platform (https://dac.institutducerveau-icm.org) were crucial to the success of this study. The mass spectrometer used for the proteomics study was funded by A.P. Møller og Hustru Chastine Mc-Kinney Møllers Fond til almene Formaal.

---

**Fig. S1:**
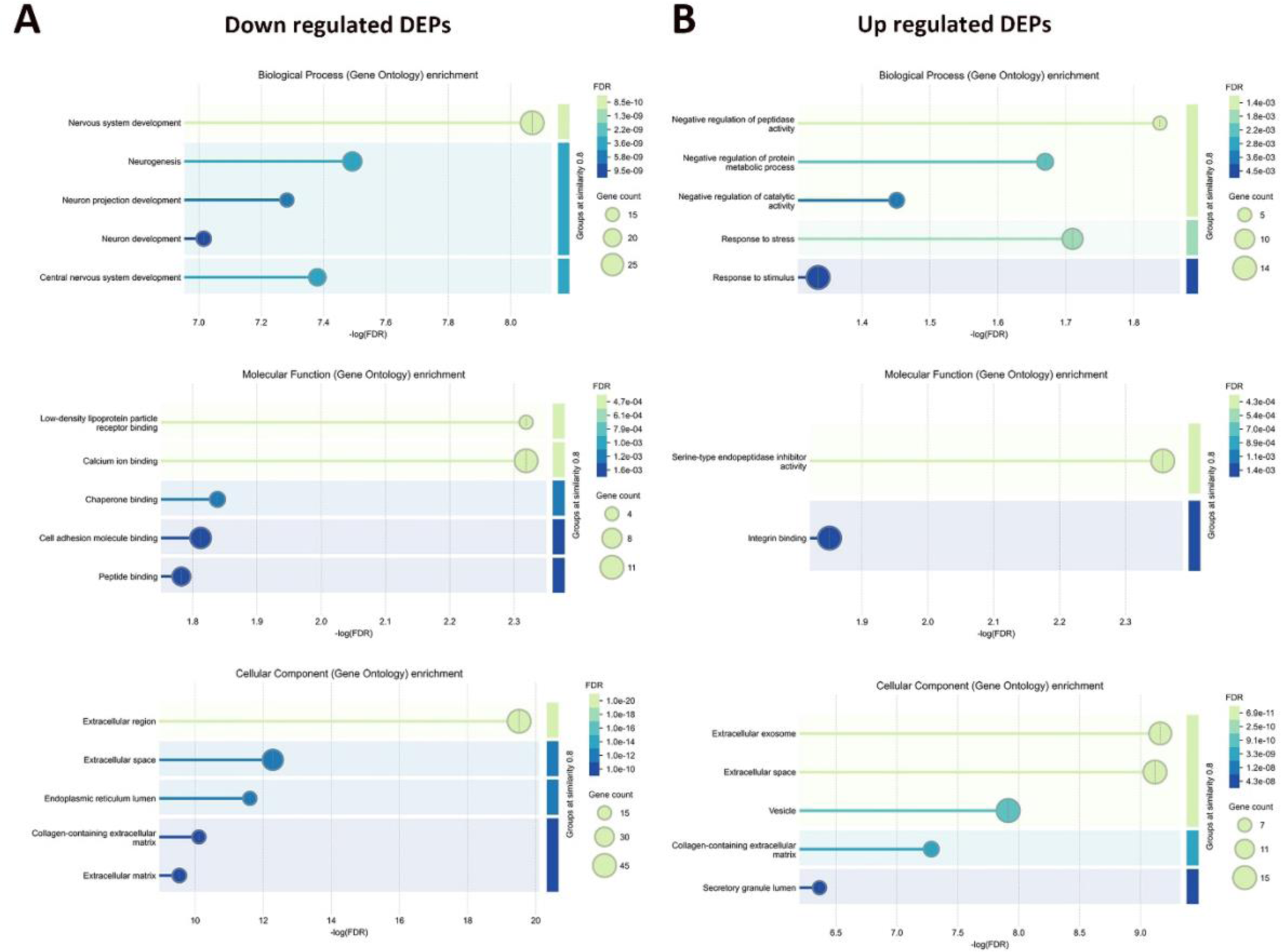
GO analysis of the DEPs. (**A**) GO analysis of the down regulated proteins; (**B**) GO analysis of up regulated proteins. According to the order of FDR, only the top 5 terms were displayed. Gene count associated to a particular GO is indicated.

**Fig. S2:**
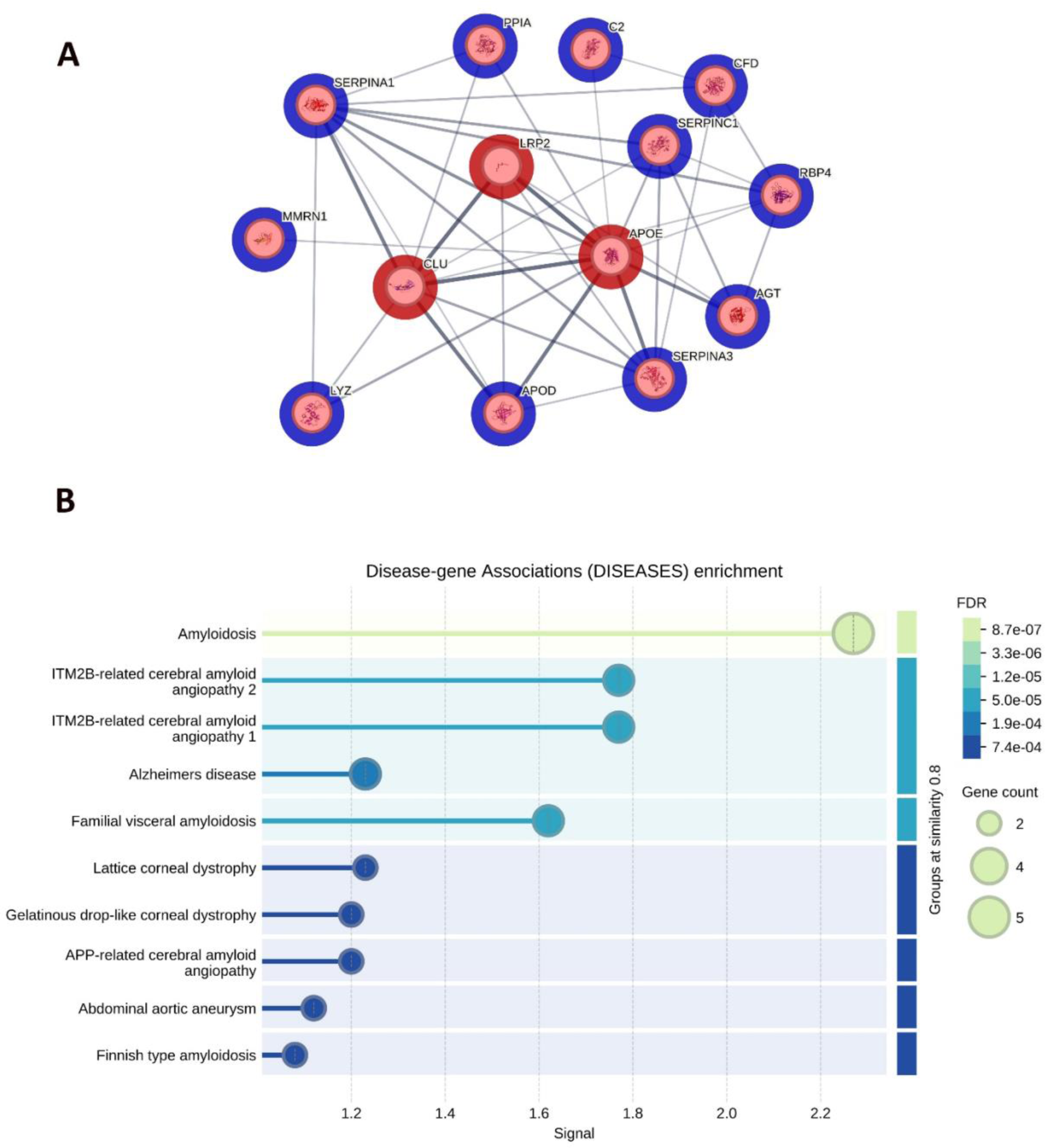
(**A**) The most significant cluster was obtained from PPI network. LRP2 and its ligands CLU and APOE were enriched as hub proteins in this cluster. (**B**) Disease-gene associations enrichment indicated an enrichment in neurodegenerative diseases.

**Fig. S3:**
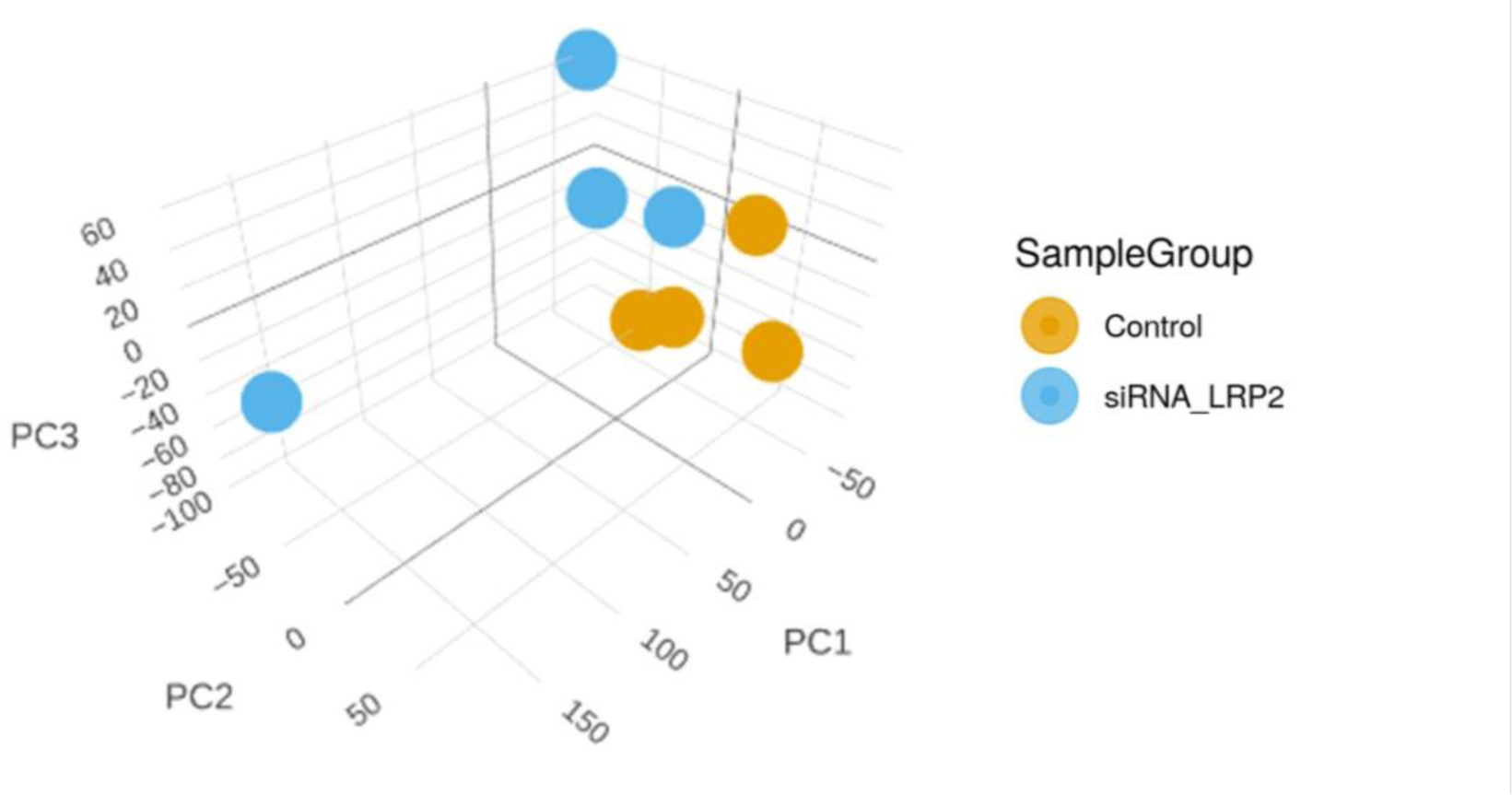
Principal component (PC) analysis, X-axis, Y-axis, and Z-axis show PC1, PC2 and PC3, respectively.

**Fig. S4:**
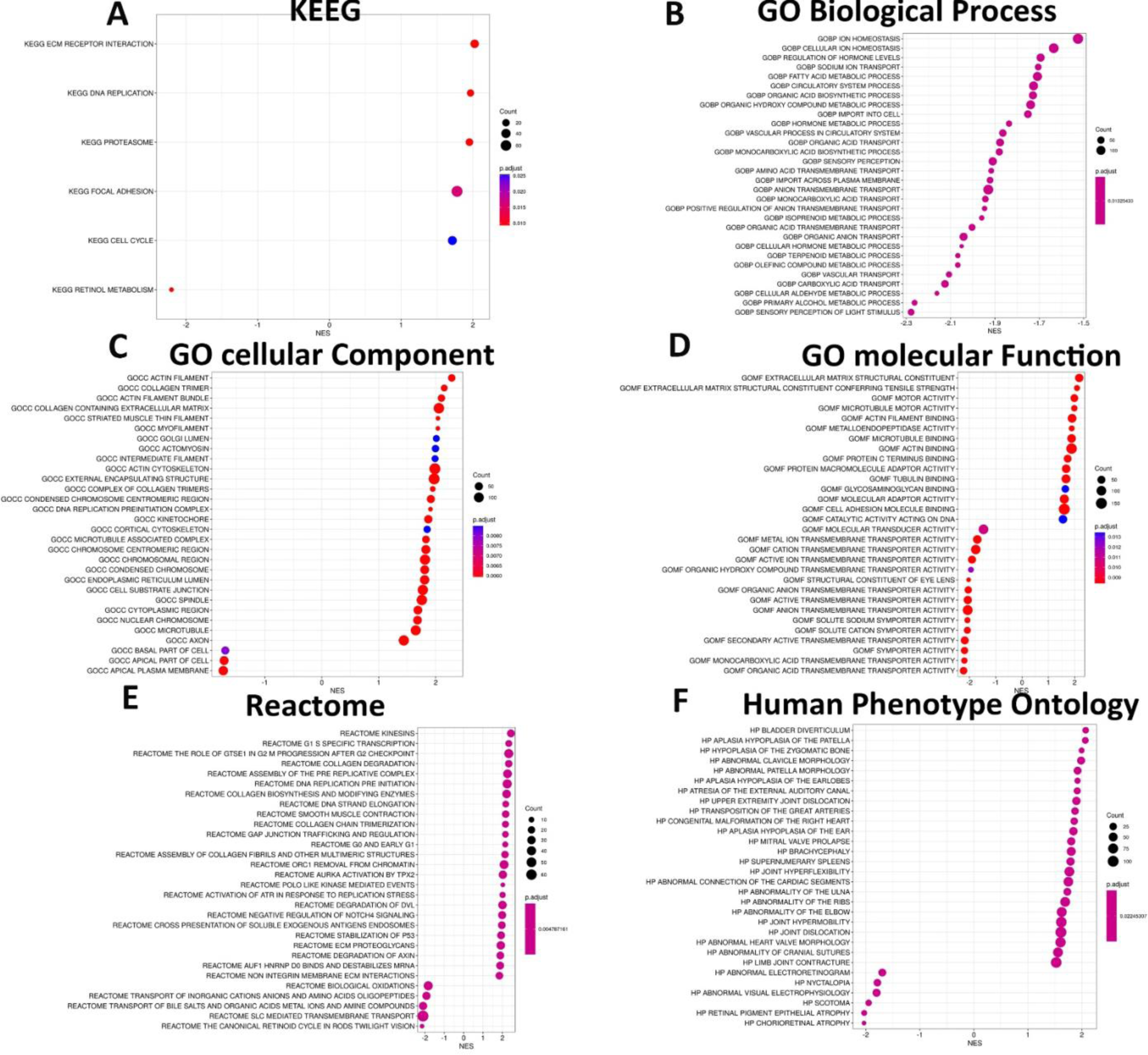
(**A**) The top 6 most enriched KEGG of downregulated DEGs and upregulated DEGs. (**B to D**) The top 30 most enriched GO terms (selected based on the p-values), (**B**) GO biological processes, (**C**) GO cellular processes, (**D**) GO molecular functions, (**E**) reactome and (**F**) human phenotype ontology. The terms were selected based on the lowest LogP values (color codes on each graph). Analysis was performed using metascape. Absolute normalized enrichment allowed to identify terms that were upregulated or downregulated using all DEGs.

**Fig. S5:**
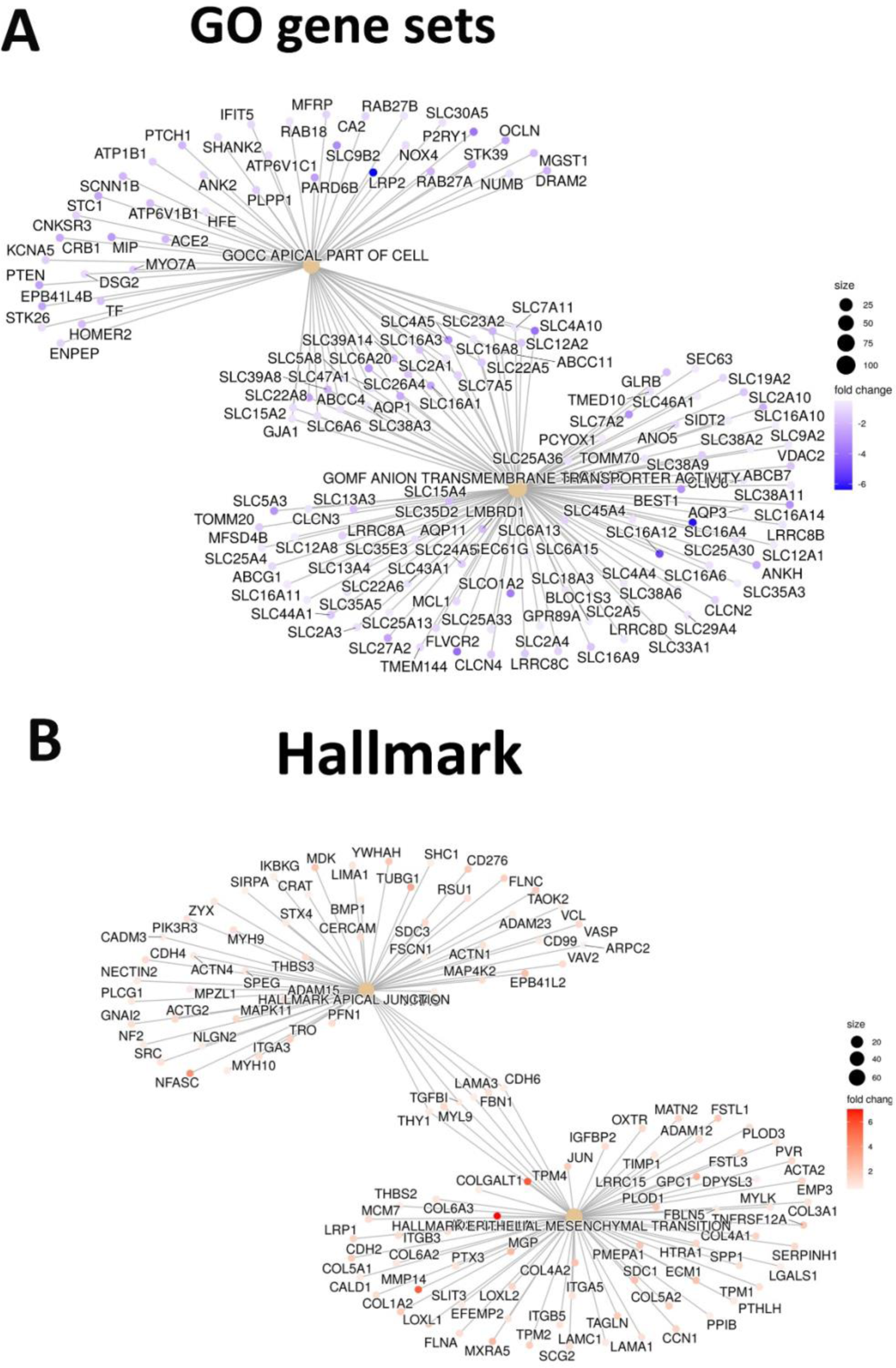
Category netplot enrichment analysis with (**A**) the top 2 most enriched GO terms (selected based on the p-values) of down-regulated DEGs and (**B**) the top 2 most enriched Hallmark of up- regulated DEGs. Analysis was performed using metascape.

